# A Transfection-Free Approach of Gene Editing via a gold-based nanoformulation of the Cas9 protein

**DOI:** 10.1101/2024.07.09.602746

**Authors:** Soultana Konstantinidou, Agnieszka Lindstaedt, Tiziana Julia Nadjeschda Schmidt, Francesco Nocilla, Giovanni Maltinti, Mafalda Angelica Rocco, Elena Landi, Alessandro De Carli, Stefania Crucitta, Michele Lai, Mauro Pistello, Valentina Cappello, Dariusz Witt, Chiara Gabellini, Piotr Barski, Vittoria Raffa

**Affiliations:** Dep. Biology, University of Pisa, SS 12 Abetone e Brennero 4, 56127, Pisa, Italy; ProChimia Surfaces Sp. z o.o., Al Zwycięstwa 96/98 F8, 81-451 Gdynia, Poland; Dep. Medicine, University of Pisa, SS 12 Abetone e Brennero 4, 56127, Pisa, Italy; Center for Materials Interfaces, Istituto Italiano di Tecnologia, Viale Rinaldo Piaggio 34, 56025 Pontedera, Italy; Dep. Clinical and Experimental Medicine, University of Pisa, via Roma 55, 56125, Pisa, Italy; Dep. Medical Biotechnologies, University of Siena, Siena, 53100, Italy; Faculty of Chemistry, Gdansk University of Technology, Narutowicza 11/12, 80-233 Gdansk, Poland

**Keywords:** CRISPR/Cas9, gold nanoparticles, gene editing, transfection-free delivery, zebrafish, human melanoma cells

## Abstract

In recent years, the CRISPR/Cas9 technology has emerged as a highly efficient tool for cell gene editing. However, the delivery of the CRISPR/Cas9 system into cells remains a significant challenge, drastically limiting *in vivo* gene therapy applications. In this study, we present a transfection/transduction-free tool for intracellular delivery of the Cas9:gRNA ribonucleoprotein. The Cas9 enzyme is conjugated to a 12 nm gold nanoparticle through affinity binding between the 6x His-tag of the protein and the NTA-Ni²LJ groups on the nanoparticles. This link chemistry allows a fine control of the density of the enzymes decorating the particle surface, the orientation of the bonding and the stability of the interaction. Importantly, the surface chemistry of this nanoformulation has been precisely engineered to modulate the cellular internalization and localization. Thanks to this approach of precision chemistry, this nanoformulation demonstrated the ability to spontaneously enter human melanoma cells as monodispersed particles that localize in cell cytoplasm, endosomes, and nucleus. It also shows effective gene editing efficiency similarly to conventional transfection tools. This gold-based formulation of Cas9 represents a ready-to-use biotech editing tool, and a promising solution for direct *in vivo* gene editing applications.

## 1. Introduction

The Clustered regularly interspaced short palindromic repeat (CRISPR) -associated protein 9 (Cas9) (CRISPR/Cas9) system has revolutionized the field of gene editing due to its simplicity and straight forward design, offering unprecedented accuracy and versatility for genome modification [1, 2]. The technology employs a ribonucleoprotein (RNP) composed of the Cas9 enzyme and a single-guide RNA (sgRNA) responsible to direct the Cas9 nuclease to a specific DNA sequence, where it induces a double-strand break (DSB) [3, 4].

Despite the advancements in biotechnology, the therapeutic application of CRISPR-Cas9 is significantly hindered by challenges regarding the efficient delivery of the editing complexes into target cells [5, 6]. Commonly, the delivery is performed via viral vectors, such as the Adeno-associated virus (AAV) and Lentivirus (LV), allowing for high transduction efficiency and for potential *in vivo* administration [6, 7]. However, the continuous expression of the system leading to high off-target events, and the limited loading capacity, immunogenicity, insertional mutagenesis and carcinogenesis posed by viral vectors lead to the increased interest for non-viral delivery systems [8]. Furthermore, viral vector manufacturing is extremely expensive, making these therapies an economic challenge for biopharmaceutical companies and national healthcare systems. The two CRISPR-based therapies - Casgevy and Lyfgenia, approved for the treatment of sickle cell disease - employ lentiviral vectors as their delivery strategy, and are priced at $2 million and $3.1 million, respectively. The high cost of these medications poses a significant barrier to accessibility, creating a major ethical issue as it raises questions about equality in healthcare. Non-viral delivery approaches, such as those using nanomedicine-based carriers, could significantly reduce these costs. Additionally, this strategy could improve the safety profile of CRISPR-based medicines through the direct administration of the RNP into cells. In fact, the RNP complex, comprising the Cas9 nuclease and the single guide RNA, is preferred due to its immediate action upon administration into the cells and limited half-life resulting in a decreased time of activity and thus mitigating possible off-target events [9, 10]. However, the RNP is a large complex, which complicates its cellular uptake, endosomal escape and stability [11]. While the mechanical methods, such as microinjection and electroporation, cannot have any *in vivo* applications due to their invasiveness [12], various nanocarriers are being adapted for a clinical setting. They have been proposed for delivery of the Cas9 RNP, such as lipid nanoparticles (LNPs), polymeric nanoparticles (PNPs), and gold nanoparticles [13, [14, 15, 16, 17, [18]. Gold nanoparticles (AuNPs), in particular, have emerged as advanced delivery platforms for gene editing due to their unique physicochemical properties and ease of functionalization [10, 18, 19]. They are exploited for directly delivering the Cas9 RNP in cells with minimal cytotoxicity due to their biocompatibility [20]. The RNP can be attached on the surface of the gold nanoparticles exploiting electrostatic interactions, as in the case of cationic arginine-gold nanoparticles (ArgNPs) interacting with negatively charged Cas9 proteins fused with an N-terminal glutamate peptide tag [21], or interactions via DNA-thiol and the RNP with the donor DNA introduced as the PAsp(DET)-coated CRISPR-Gold nanoparticles [22]. Aiming at controlled release by endosomal disruption and targeted delivery, gold nanoparticles can also be functionalized with various ligands and programmed for response in internal stimuli, including pH, enzymes, ATP, redox, and oxygen levels [23, 24, 25].

In this study, we designed a gold-based nanoformulation with highly predictable performance to meet future regulatory standards for application in the genome editing field. We used a functionalization scheme that allows fine control of the number of enzyme molecules per particle and a highly oriented and reproducible bonding between the carrier and its cargo.

Specifically, the Cas9 enzyme was conjugated via the 6x His tag to a 12 nm spherical gold nanoparticle (AuNP) terminated with nitrilotriacetic acid (NTA) groups. The background chemistry of the AuNP was precisely designed to modulate cell internalization, trafficking, and localization of the AuNP-Cas9:gRNA formulation. *In vitro* and *in vivo* experiments demonstrate the catalytic activity of the AuNP-Cas9, highlighting its gene-editing efficiency in both zebrafish (*Danio rerio*) and human melanoma cells (A375). This novel AuNP-Cas9 formulation is a robust and versatile platform that enables the Cas9 enzyme to effectively internalize spontaneously and cleave target DNA within human cells without the need for conventional transfection or transduction methods, offering a significant advancement in the delivery and application of CRISPR-Cas9 technology.

## 2. Results and Discussion

### 2.1. Synthesis and physical characterization of the AuNP-Cas9

Gold nanoparticles (AuNPs) with 12 nm size were prepared and stabilized with polyethylene thiol terminated (AMW 3kDa) with carboxylic group or amino groups. The particles terminating with carboxylic groups were activated by formation of active esters (sulfoNHS) and treated with thiolated NTA (AuNP.NTA.C). The activation of carboxylic groups did not consume all available carboxylic groups and the AuNP.NTA.C possessed some free carboxylic acid groups to provide appropriate background and improved solubility in aqueous solution. The AuNP.NTA.C had an inorganic core size of 11 ± 3nm, hydrodynamic size of 26 ± 6nm and Zeta potential of -35 ± 5mV (see supplementary data Figure S1). Subsequent treatment with nickel (Ni^2+^) salt and the Cas9 with His tag allowed attachment of the protein (Figure 1A).

**Figure 1.**
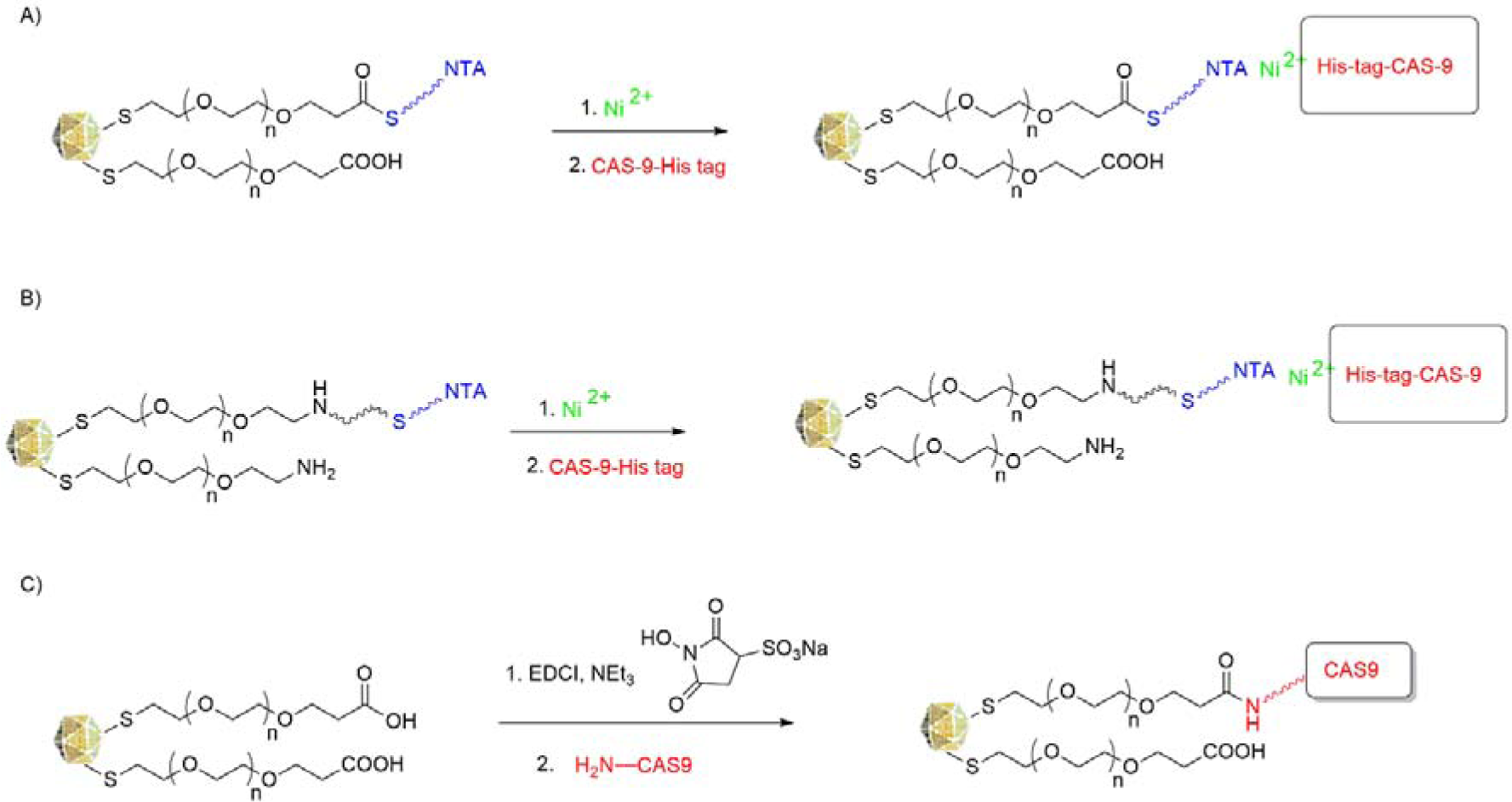
Preparation of functionalized gold nanoparticles with Cas9. (A) Gold nanoparticles with amino background and NTA. (B) Gold nanoparticles with carboxylic background and NTA. Polyethylene glycol thiol terminated with an amino and carboxyl group (average MW 3kDa) were used respectively. (C) Gold nanoparticles with carboxylic background with NHS.

For particles terminating with amino groups, AuNPs with 12 nm size were prepared and stabilized with polyethylene thiol terminated with amino group (AMW 3kDa). After purification, some of the terminal amino groups were treated with active ester with maleimide functionality. This functionality allowed efficient attachment of NTA by the reaction of maleimide functionality with NTA terminated thiol (AuNP.NTA.N). This functionalization did not consume all available amino groups. The amount of NTA moieties on the surface was calculated by analyzing the complexed nickel ions via Microwave-Induced Plasma - Optical Emission Spectrometry (MIP-OES), resulting in 4% (NTA terminated groups / amino groups). Since the result obtained with the MIP-OES method falls into the limit of detection range, we decided to further support it with the Inductively Coupled Plasma Spectroscopy (ICP) method and the amount of NTA groups on the total of the amino terminating groups was 6%. The final AuNP.NTA.N possessed an inorganic core size of 11 ± 3nm, a hydrodynamic size of 23 ± 5nm and a Z-potential of -2 ± 20mV (see supplementary data Figure S2). Similarly, subsequent treatment with Ni^2+^ salt allowed attachment of the Cas9 protein via the His tag (Figure 1B).

Finally, only for comparative purpose, we used a conventional NHS chemistry. AuNPs stabilized with polyethylene thiol terminated with carboxylic group were activated by formation of active esters (sulfoNHS) and treated with amino-terminated Cas9 (Figure 1C).

Cas9 protein binding was firstly confirmed by SDS Page (Figure S3). In order to determine a more precise concentration of the Cas9 bound protein onto the surface of the nanoparticles, we performed a Dot Blot assay (Figure S4). For AuNP.NTA.N, the binding efficiency of Cas9 was 1.2±0.26 ng of Cas9 per 10^8^ AuNP (mean±SEM, n=11, single data available in Table S1). Interestingly, the theoretical calculations of Cas9 binding ability based on ICP analysis (6% of NTA groups onto the particle surface) was corresponding to 50 molecules Cas9 molecules per AuNP nanoparticles. This value perfectly matches the quantification of Cas9 protein by Dot Blot data, consisting of 45.1±9.6 molecules of Cas9 per AuNP (mean±SEM, n=11, single data available in Table S1), confirming the robustness of the proposed functionalization scheme.

Three different batches of AuNP.NTA.C and AuNP.NTA.N were analysed 1,3 and 6 months after synthesis to define the shelf-life. Particle degradation can be easily estimated on the basis of the drift over the time from typical values of the surface plasmon resonance (SPR) maximum peak, size (by DLS) and zeta potential. The characteristic data are presented in Table 1, documenting that AuNP.NTA.C and AuNP.NTA.N are stable for more than 6 months when stored in a pure state in the temperature range of 20-30°C (preferably at RT) and dispersed in deionized water or 20mM TAPS pH 8.0, respectively.

**Table 1.** Characteristic data for AuNP.NTA.C and AuNP.NTA.N at 1, 3, 6 months after the synthesis (n=3).

|  | AuNP.NTA.C |  |  |  | AuNP.NTA.N |  |  |  |
| --- | --- | --- | --- | --- | --- | --- | --- | --- |
|  | 0 month | 1 month | 3 months | 6 months | 0 month | 1 month | 3 months | 6 months |
| Size (DLS) | 26 nm ± 5 nm | 25 nm ± 5 nm | 26 nm ± 6 nm | 26 nm ± 5 nm | 22 nm ± 5 nm | 23 nm ± 5 nm | 23 nm ± 5 nm | 24 nm ± 5 nm |
| SPR max | 523 nm ± 2 nm | 522 nm ± 3 nm | 522 nm ± 2 nm | 522 ± 2 nm | 523 nm ± 2 nm | 523 nm ± 2 nm | 524 nm ± 3 nm | 523 nm ± 3 nm |
| Zeta potential | -35 mV ± 3 mV | -35 mV ± 3 mV | -34 mV ± 3 mV | -31 mV ± 5 nm | -7 mV ± 8 mV | -2 mV ± 6 mV | -3 mV ± 7 mV | -5 mV ± 6 nm |

We also estimated that the Cas9 protein does not detach from functionalised AuNP.NTA.C- Cas9 and AuNP.NTA.N-Cas9 up to six weeks, when they are stored in a pure state at -20°C in 50% glycerol (Figure S5 and S6).

### 2.2. In Vitro characterization of the AuNP-Cas9

The catalytic activity of the various nanoformulations of the Cas9 *in vitro* was tested on a linear DNA fragment including the target sequence of the gRNA (Figure 2). While no DNA cleavage was observed by the AuNP.NHS-Cas9:gRNA (Figure 2A), both the AuNP.NTA.C- Cas9:gRNA and AuNP.NTA.N-Cas9:gRNA were able to recognize and cut the target DNA in a dose-dependent way after the functionalization process (Figure 2B, C). However, the AuNP.NTA.N-Cas9:gRNA worked more efficiently than the AuNP.NTA.C-Cas9:gRNA in all the tested concentrations, (p<0.0001 for 150 nM and 100 nM, and p=0.0063 for 50 nM, Figure 2D). Indeed, there was no statistical difference in the cleavage efficiency using the two highest investigated doses (150 nM and 100 nM) between the AuNP.NTA.N-Cas9:gRNA and the non-conjugated Cas9:gRNA, p=0.9779 and p=0.1762, respectively (Figure 2D).

**Figure 2.**
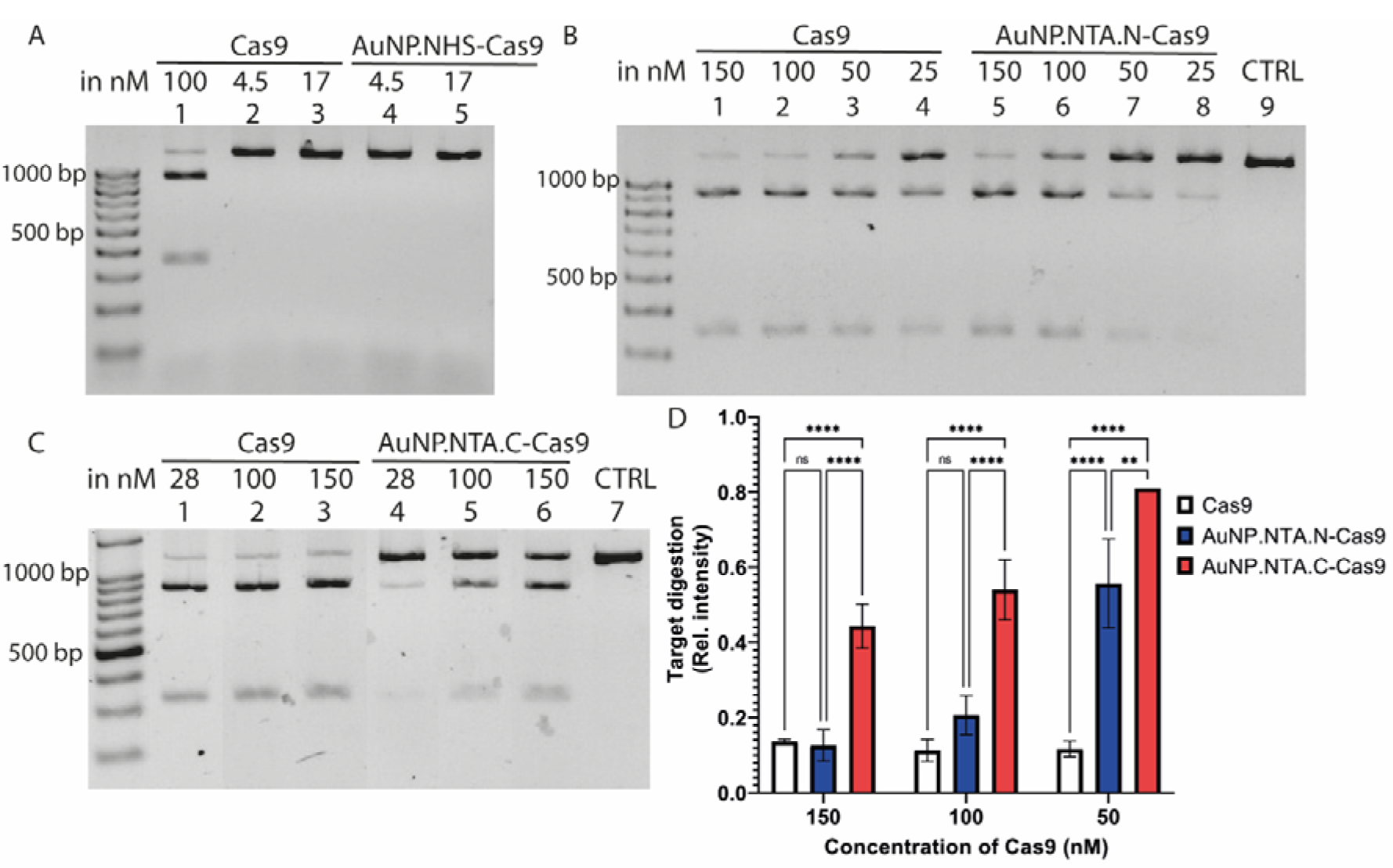
Endonuclease activity of the AuNP-Cas9. (A) AuNP-Cas9 consisted of AuNP.NHS nanoparticle, (B) AuNP-Cas9 with AuNP.NTA.N nanoparticle, (C) AuNP-Cas9 with AuNP.NTA.C nanoparticle. (D) Quantification of endonuclease efficiency of the AuNP-Cas9 nanoformulations, expressed as relative band intensity (A-D) Sample including non-conjugated Cas9 was used as positive control.. Statistical analysis was performed by 2-way ANOVA (Tukey’s multiple comparisons test), Row Factor (dose): p<0.0001; Column Factor (treatment): p<0.0001. For 150 nM: comparison between Cas9 and AuNP.NTA.N-Cas9 p=0.9779, between Cas9 and AuNP.NTA.C-Cas9 p<0.0001, and between AuNP.NTA.N- Cas9 and AuNP.NTA.C-Cas9 p<0.0001. For 100 nM: comparison between Cas9 and AuNP.NTA.N-Cas9 p=0.1762, between Cas9 and AuNP.NTA.C-Cas9 p<0.0001, and between AuNP.NTA.N-Cas9 and AuNP.NTA.C-Cas9 p<0.0001. For 50 nM: comparison between Cas9 and AuNP.NTA.N-Cas9 p<0.0001, between Cas9 and AuNP.NTA.C-Cas9 p<0.0001, and between AuNP.NTA.N-Cas9 and AuNP.NTA.C-Cas9 p=0.0063. N=3 for each condition.

The nanoparticles were further characterized for their biocompatibility in human melanoma A375 cells. The AuNP.NHS nanoparticles showed a metabolic impact in a dose-response manner (Figure S7A), with a strong correlation (one-phase decay, R²=0.99) and an inhibition concentration (IC25) of 7.5 x 10¹¹ NPs/ml. This effect may be attributed to the presence of NHS active esters on the nanoparticle surface, which can interact with cytosolic molecules. In contrast, AuNP.NTA nanoparticles (with both backgrounds) did not affect the metabolic activity of A375 cells at the tested concentrations (Figure S7B-C). These results concluded the non-suitability of the NHS nanoparticles.

Then, the cellular internalization and nuclear localization of the nanoformulations were investigated in human melanoma A375 line. Cells were incubated with the AuNP.NTA.N- Cas9:gRNA or AuNP.NTA.C-Cas9:gRNA without any transfection reagent (Figure 3A). As demonstrated by live-cell imaging, the AuNP.NTA.C-Cas9 enters the cells and the nucleus after 1 hour of incubation (Supplementary information). Since the Cas9:gRNA RNP is frequently delivered into cells by lipofection, we used RNAiMAX kit for delivery of the RNP, as positive control. Both the nanoformulations with the different backgrounds were able to enter in the nucleus of the A375 cells after 2 hours of incubation (Figure 3B-C). The nuclear fluorescence intensity signal of the Cas9 for AuNP.NTA.C-Cas9:gRNA is not statistically different to the one of the lipofection method (p=0.1411), while AuNP.NTA.N-Cas9 shows a stronger nuclear fluorescence intensity compared to the lipofection method (p<0.0001) (Figure 3D-E).

**Figure 3.**
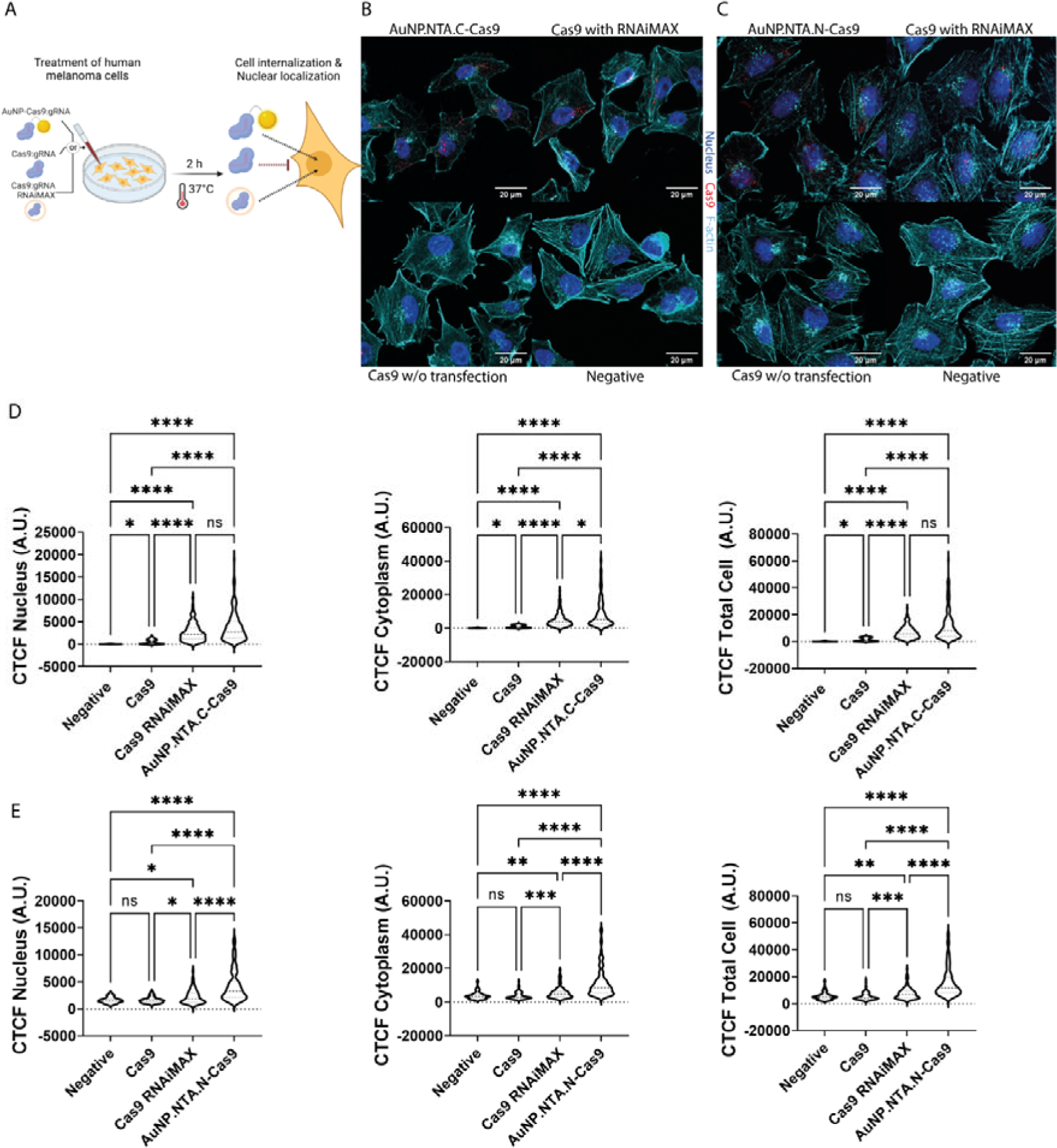
Internalization and nuclear localization of the AuNP-Cas9 in A375 cells. (A) Schematic illustration of the experimental protocol. Representative confocal image of A375 cells treated with (B) AuNP.NTA.C-Cas9:gRNA and (C) AuNP.NTA.N-Cas9:gRNA. (D and E) Violin plots representing the Cas9 fluorescence intensity as correct total cell fluorescence (CTCF) detected in the nucleus, cytoplasm and total cell. B-E) Cells exposed to Cas9:gRNA with and without RNAiMAX were used as positive and negative controls, respectively.

Statistical analysis was performed by the Kruskal-Wallis test, p<0.0001. (D) Negative control (N= 58); Cas9:gRNA (N=54); Cas9:gRNA RNAiMAX (N=152); AuNP.NTA.C-Cas9:gRNA (N=133). (E) Negative control (N= 98); Cas9:gRNA (N=108); Cas9:gRNA RNAiMAX (N=98); AuNP.NTA.N-Cas9:gRNA (N=195).

Next, we detected AuNP localization via transmission electron microscopy (Figure 4A). In cells incubated with AuNP.NTA.C-Cas9:gRNA, particles were detected in an aggregated form (Figure 4B). In cells incubated with AuNP.NTA.N-Cas9:gRNA, particles were detected in the cytoplasm of A375 cells after 30 minutes or 1 hour of incubation with the cells, without any transfection reagent (Figure S8). After 5 hours of incubation with the A375 cells, non-functionalized nanoparticles (AuNP.NTA.N) were found inside A375 cells mostly close to the cell membrane and in agglomeration forms (Figure S9). In cells incubated with the functionalised ones (AuNP.NTA.N-Cas9:gRNA), particles entered the cells without any transfection method and they were localised in endosomes, lysosomes, as well as in the cytoplasm and nucleus always in a monodispersed form (Figure 4C).

**Figure 4.**
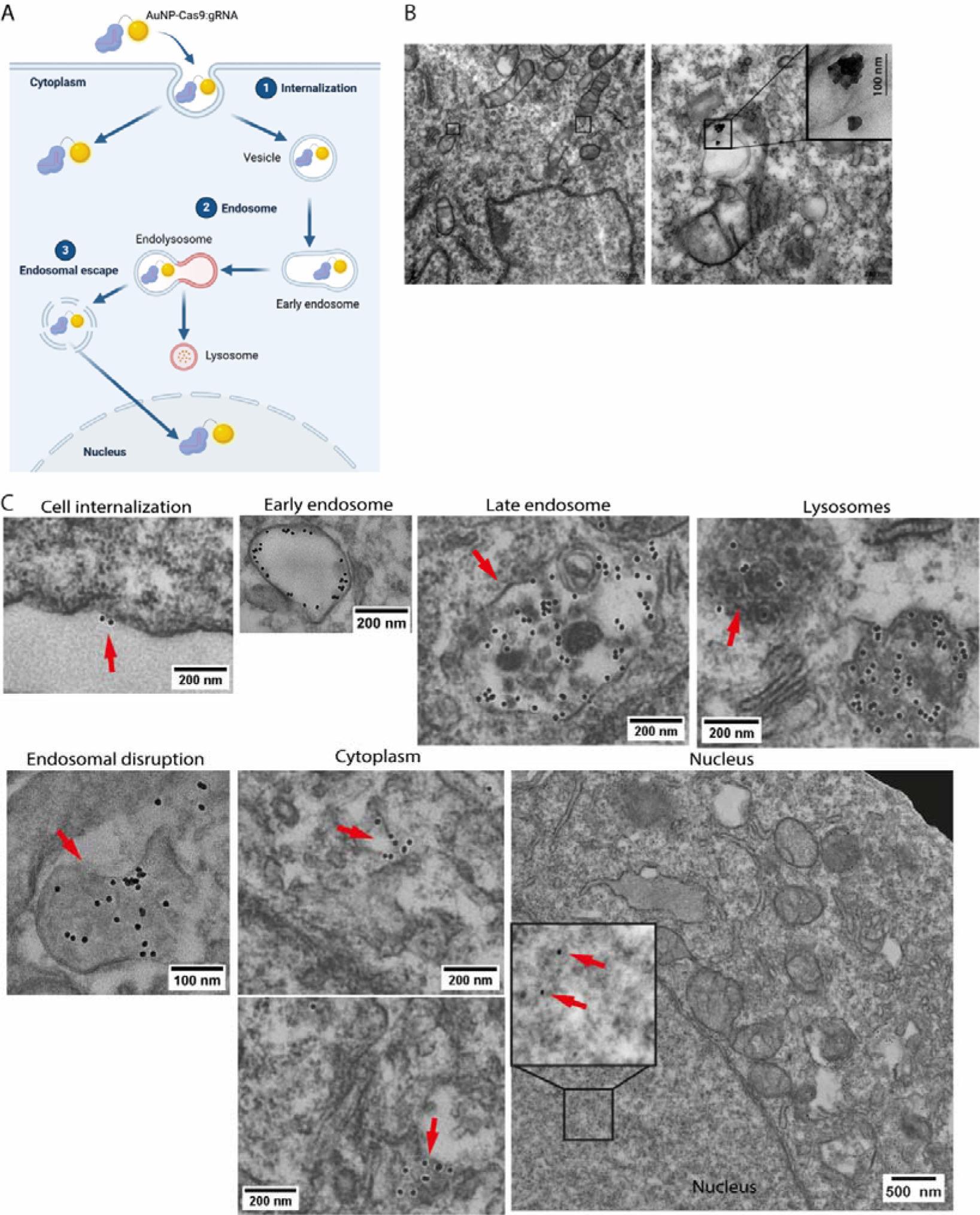
Intracellular localization of the AuNP-Cas9 in human melanoma A375 cells. (A) Illustration of the intracellular detection of the gold nanoformulation. (B) TEM images of human melanoma cells A375 incubated with the AuNP.NTA.C-Cas9:gRNA for 2 hours. The scale bar for the large images is 500 nm and for the magnification 100 nm. (C) TEM images of human melanoma cells A375 incubated with the AuNP.NTA.N-Cas9:gRNA for 5 hours. The scale bars are presented in each image.

Taken all together, it was concluded that the amino background on the nanoparticles is preferred for the synthesis of the nanoformulation and further experiments were conducted using the AuNP.NTA.N-Cas9:gRNA, simply named from now on AuNP-Cas9:gRNA.

### 2.3. Gene editing Activity of the AuNP-Cas9 in human melanoma cells and in vivo, in zebrafish

#### 2.3.1 AuNP-Cas9 edited ASAH1 gene in A375 melanoma cells

With the aim to probe the editing efficiency of the AuNP-Cas9, we administered the nanoformulation directly into cell media of A375 cells, using as a reference control the cells lipofected with Cas9 RNPs. Briefly, the AuNP-Cas9 was administered at concentrations of 10 and 55.5 nM, then extracted genomic DNA from the cells after 48 h to perform digital droplet PCR (ddPCR), as schematically represented in Figure 5A. To highlight the editing, we designed two probes targeting the editing region of the ASAH1 gene. The first one (HEX) recognizes the same nucleotides targeted by the RNPs, while the second (FAM) recognizes a region downstream that is not targeted by RNPs (Figure 5A). In this way, all samples must be positive for the FAM signal, while if gene editing occurs, the HEX signal is lost. As shown in Figure 5B, AuNP-Cas9 administered at 10 nM had the same, even if lower, editing efficiency of lipofectamine-delivered RNPs. Moreover, AuNP-Cas9 delivered at 55.5 nM increased the editing efficiency to 1.2%, showing a dose-dependent activity oft he nanoformulation.

**Figure 5.**
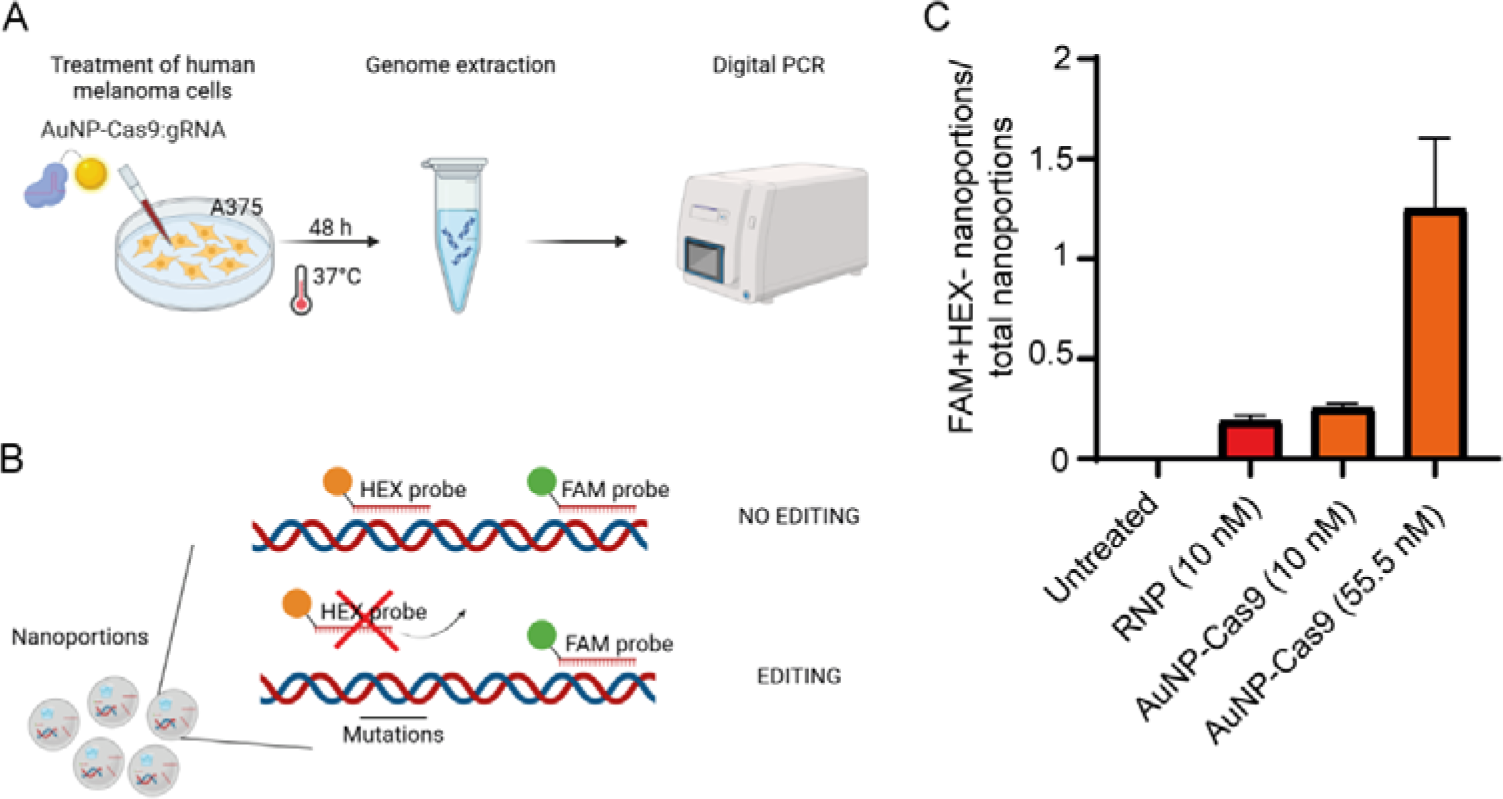
AuNP-Cas9 edited target ASAH1 gene in A375 melanoma cells. (A) Representative experimental process scheme. Briefly we treated A375 cells with the AuNP- Cas9, then genomic DNA was extracted and probed by dPCR. (B) Digital PCR set up and read-out illustration. (C) FAM+HEX- nanopartitions out of total.

#### 2.3.2 AuNP-Cas9 edited the tyrosinase gene in zebrafish

Furthermore, we evaluated the gene editing efficiency *in vivo*, using zebrafish (Danio rerio) asanimal model,. We selected to target the tyrosinase (tyr) gene, encoding for an enzyme required for the synthesis of eumelanin, the pigment contained in zebrafish melanocytes. Loss of function of the tyr gene results in albino-like phenotype of zebrafish embryos, that lack melanin pigmentation in the eye area and in the body [26, [27]. Zebrafish embryos were injected with the AuNP-Cas9:gRNA in the zygote and their development was checked till 3 days post fertilization (dpf) (Figure 6A). When the nanoformulation of Cas9 in complex with the different gRNAs tyr1, tyr2 or tyr3 (AuNP-Cas9:gRNA) was injected, it seemed to affect the viability of the embryos (p=0.0170, p=0.0190 and p=0.1262, respectively), but the survival ratio towards the wild-type embryos was for all the cases more than 0.8, thus, it was not considered significant (Figure 6B). In addition, injection of the unconjugated Cas9:gRNA tyr2 in a concentration of 1000 pg Cas9/embryo affected the viability of the embryos, p<0.0001, while the concentration of 750 pg Cas9/embryo did not present a significant mortality ratio, p=0.2208 (Figure S10), concluding that the presence of the Cas9 protein in high concentrations can impact embryo survivals.

**Figure 6.**
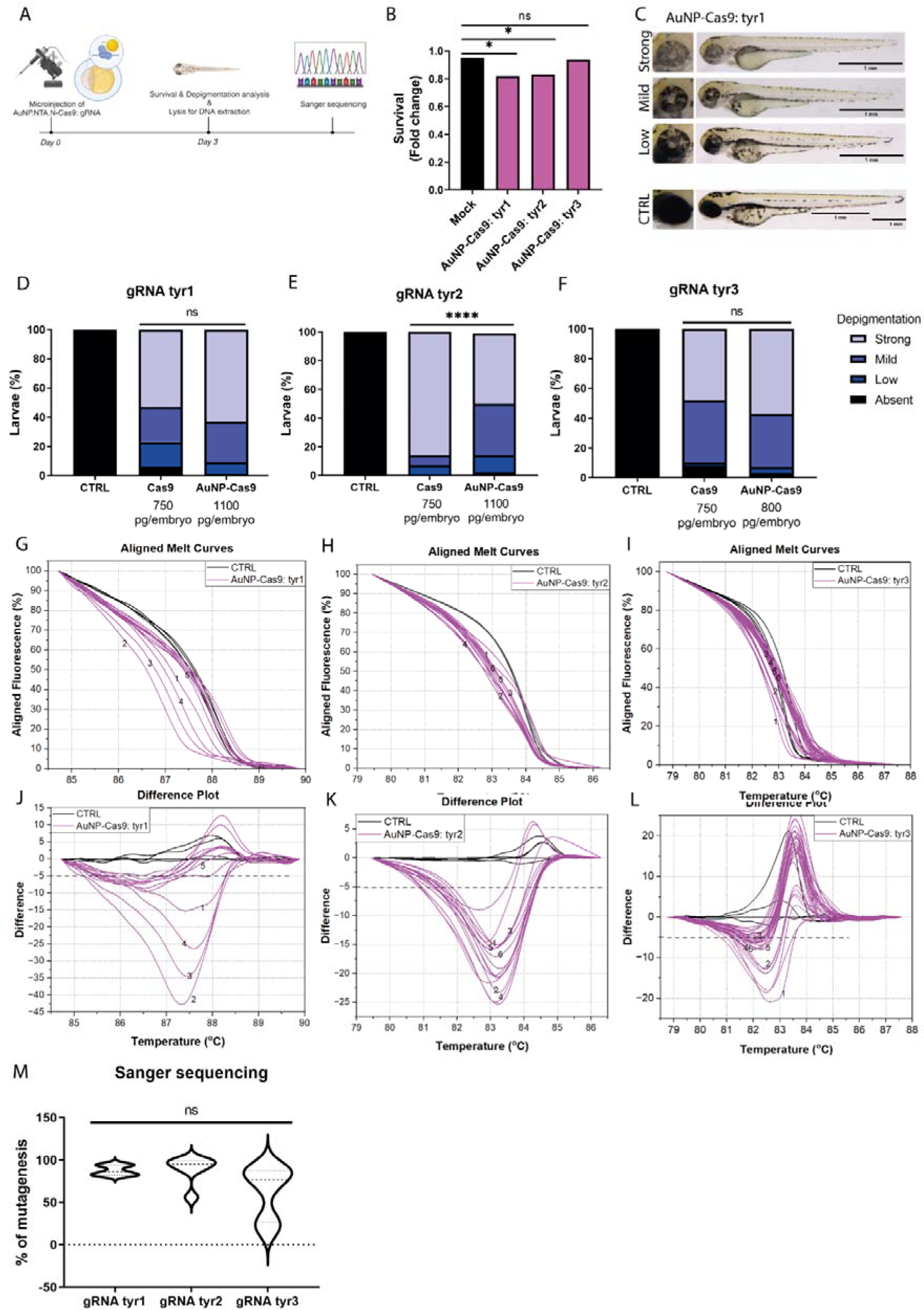
Gene editing efficiency of the AuNP-Cas9 *in vivo*. (A) Schematic representation of zebrafish embryo microinjection with nanoformulations targeting tyrosinase (tyr) gene and evaluation of gene editing by HRM assay. (B) Toxicity analysis of the AuNP-Cas9:gRNA. A one-sided χ2 test was performed and every group was analysed towards the control of the same experiment. Data are shown as survival ratio of every treatment normalised towards the control of the same experiment; for AuNP-Cas9:tyr1 N=125, p value=0.0170; for AuNP- Cas9:tyr2 N=53, p value=0.0190, for AuNP-Cas9:tyr3 N=142, p value=0.1262. (C) Representative images of zebrafish larvae injected with AuNP-Cas9:tyr1 . Wild-type zebrafish have the same phenotype as the group “Absent” (images not shown). (D) Quantification of depigmented embryos injected with gRNA tyr1Statistical analysis was done by χ2 test, p=0.2833. For Cas9:tyr1 N=86, and AuNP-Cas9:tyr1 N=72. (E) Quantification of depigmented embryos injected with gRNA tyr2.Statistical analysis was done by χ2 test, p<0.0001. For Cas9:tyr2 N=107, and AuNP-Cas9:tyr2 N=36. (F) Quantification of depigmented embryos injected with gRNA tyr3. Statistical analysis was done by χ2 test, p=0.5161. For Cas9:tyr3 N=89, and AuNP-Cas9:tyr3 N=56. (G-L) Melting analysis performed by High Resolution Melt Software (Applied Biosystems). The aligned melt curves and difference plots are shown for each treatment. In the Difference Plots, an uninjected control sample was selected as a reference (baseline=0). The control for all the panels is wild-type embryos. (M) Sanger sequencing analysis. The percentage of mutagenesis for the treatment AuNP-Cas9: gRNA tyr1 or tyr2 or tyr3 are shown. Statistical analysis was performed by Kruskal-Wallis test, p=0.9941, p=7419, p=0.0768, respectively. The microinjections of the zebrafish were performed in duplicate.

The injected zebrafish presented three different types of depigmentation at 3dpf: strong, mild or low (Figure 6C). Interestingly, the caused phenotypic effect was similar with the depigmentation phenotype derived by the unconjugated Cas9 for the gRNAs tyr1 and tyr3 (p=0.2833, p=0.5161, respectively), but using the tyr2 there was a statistical difference, p<0.0001 (Figure 6D-F). Moreover, the DNA of 3dpf larvae was extracted and analysed for modifications and indel mutations by High Resolution Melting (HRM) and Sanger sequencing. The presence of modifications/mutations in the region of interest lead to the formation of mosaic embryos presenting different DNA melting temperature profiles. Indeed, the target DNA region of zebrafish injected with the AuNP-Cas9 in complex with the different gRNAs presented different melting profiles than the control non-injected embryos, as demonstrated in the graphs titled “Aligned Melt Curves” and “Difference Plot” (Figure 6G- L), where the melting profile of a control zebrafish was selected as a reference. Sanger sequencing analysis revealed the presence of indel mutations in the region of interest (Figure 6M, Figure S11). In specific, the gRNA tyr1 and tyr3 induced a variety of deletions and the gRNA tyr2 mostly a deletion of 4 nucleotides. The induced percentage of mutagenesis was similar for all the treatments irrelevantly from the used gRNA (Figure 6M). All the data from the sequencing analysis by the ICE software are provided as supporting information (Figure S11).

## 3. Conclusion

In this study, we generated a gold-based formulation of the Cas9:gRNA ribonucleoprotein. Since a precise link chemistry is essential for not compromising the translocase, helicase and nuclease activity of Cas9 protein, we opted for a link chemistry that immobilizes the enzyme with a precise orientation onto the nanoparticle surface. Since the mobility of the enzyme is equally important for its activity, we used polyethylene glycol spacers. The spacer was amino terminated to facilitate the internalization of the nanoformulation as monodispersed nanoparticles. Additionally, the density of the NTA tag was optimized at the value of the 6% of the terminating groups - corresponding to 50 molecules of enzyme per particle - to avoid loss of activity due to steric hindrance. As a result of this approach of precision chemistry, this nanoformulation demonstrated a cleavage efficiency similarly to free Cas9:gRNA. The AuNP-Cas9:gRNA is very stable and can be stored for 6 weeks at -20C in 50% glycerol. It is not cytotoxic or genotoxic, as documented by experiments performed in human melanoma cells A375 and zebrafish embryos. AuNP-Cas9:gRNA can spontaneously enter the cells without any transfection tool and the internalisation efficiency is not statistically different from that obtained using a standard lipofection kit for transfecting Cas proteins in eukaryotic cells. When A375 cells are incubated with the AuNP-Cas9:gRNA, the ribonucleoprotein show nuclear localization, with a peak in 1-2 hours after the administration. The nanoparticles have been found to localise in cell cytoplasm, nucleus and endosomes. AuNP-Cas9:gRNA injected in zebrafish embryos showed the ability to perform gene editing, as documented by experimental data at the phenotypic level (depigmented phenotype) and at the molecular level (HRM assay, Sanger sequencing). The AuNP-Cas9:gRNA has also the capability to bind to the target and to perform gene editing when incubated with human melanoma cells, as documented by digital droplet PCR (ddPCR).

## 4. Experimental Section/Methods

### Preparation of Gold nanoparticles

Gold nanoparticles (AuNPs, 12 nm) stabilized with citrate were prepared according to a Turkevich method [28]. Trisodium citrate (Sigma-Aldrich, USA) was used for the reduction of gold chloride trihydrate solution (HAuCl_4,_ Thermo Fisher Scientific, Belgium) in aqueous solution near boiling point.

### Preparation of Gold nanoparticles with NTA and carboxylic background

AuNPs with 12 nm size were prepared and stabilized with polyethylene thiol terminated with carboxylic group (average MW 3kDa, ProChimia Surfaces, Poland). After purification, terminal carboxylic groups were activated by formation of active esters (sulfo-NHS, Sigma-Aldrich, Germany) and treated with thiolated NTA (ProChimia Surfaces, Poland). Size, zeta potential and spectroscopic features were under investigation at each step of the synthesis.

### Preparation of Gold nanoparticles with NTA amino background

AuNPs with 12 nm size were prepared and stabilized with polyethylene thiol terminated with amino group (average MW 3kDa, ProChimia Surfaces, Poland). After purification, some of the terminal amino groups were treated with active ester with maleimide functionality (ProChimia Surfaces, Poland). This functionality allowed efficient attachment of NTA by the reaction of maleimide functionality with NTA terminated thiol (ProChimia Surfaces, Poland). Size, zeta potential and spectroscopic features were under investigation at each step of the synthesis.

### Preparation of Gold nanoparticles with NHS and carboxylic background

AuNPs with 12 nm size were prepared and stabilized with polyethylene thiol terminated with carboxylic group (average MW 3kDa, ProChimia Surfaces, Poland). After purification, some of the terminal carboxylic groups were treated with sulfo-NHS (Sigma-Aldrich, Germany) and 1-ethyl-3-carbodiimide hydrochloride (EDCI, Sigma-Aldrich, Germany)) with triethylamine (NEt3, Sigma-Aldrich, Germany).

### Protein binding

Subsequent treatment with nickel (Ni^2+^) salt and Cas9 with His tag allowed attachment of Cas9 for the generation of the NP. The activation of carboxylic or amino groups and attachment of Cas9 did not consume all available terminal groups. The final AuNP-Cas9 complex possessed some free carboxylic or amino groups to provide appropriate background and improved solubility in aqueous solution.

### Analytical methods for AuNP quality control

The spectroscopic properties of the AuNP were characterised by UV-Vis spectroscopy. The synthesised AuNPs exhibit the typical optical absorbance band in the visible region (500 nm- 600 nm) due to the known localised surface plasmon resonance. UV-Vis analysis also allows to determine the concentration of obtained AuNPs after synthesis. Spectral analysis was performed with Lambda 365 (PerkinElmer Fisher Scientific, USA) double beam diode array spectrometer to collect spectra from 300–700 nm using a slit width of 1 nm. Spectra were collected for 1ml samples using a cell with a path length of 1 cm.

For the determination of the size of the obtained AuNPs, Dynamic light scattering (DLS) and Transmission electron microscopy (TEM) were used. These methods of AuNPs characterization are necessary to determine the degree of polydispersity of the nanoparticles relevant for product quality. Hydrodynamic diameters values were measured with Zetasizer Ultra (Malvern Panalytical Ltd, UK) for 50mM solutions of AuNPs in 13° and 173° backscatter mode. For AuNPs imaging we used a JEOL 1010 Transmission Electron Microscope operating at an accelerating voltage of 100 keV and an AMT XR41-B 4- megapixel (2048) bottom mount CCD camera.

Zeta Potential analysis was used to determine if a surface modification of the nanoparticle has been successful or if the processing step has modified the nanoparticle surface. All measurements of zeta potentials were performed using a ZetaSizer Ultra system (Malvern Panalytical Ltd, UK) with 50 nM solutions of AuNPs.

Each AuNP construct was initially analysed for its ligands theoretical lengths and functional corona orientation of each ligand to estimate the maximum size. These values were compared versus hydrodynamic diameter DLS measurements and metallic core size calculated by TEM analysis.

Nickel analysis was performed utilising Microwave-Induced Plasma - Optical Emission Spectrometry (MIP-OES) employing Agilent’s MP-AES 4210 instrument. The determination of nickel involved different wavelengths (352.450 nm and 361.940 nm), with the calculated arithmetic mean considered as the final analysis result. Accordingly, nickel determinations using the Inductively Coupled Plasma Spectroscopy (ICP) method were performed using the ICP-OES 5800 spectrometer (Agilent) equipped with a cyclone fog chamber and a pneumatic sprayer. Yttrium was used as an internal standard.

### Dot blot assay

Quantification of bound protein on the surface of the Au NPs was performed by Dot Blot assay on nitrocellulose membrane (Amersham™ Protran™ Premium 0,45 μm NC, nitrocellulose blotting membrane, GE Healthcare Life Science, Cod 10600003). 2μl of sample were spotted and compared with known concentrations of Cas9 (Alt-R S.p. HiFi Cas9 Nuclease V3) (40 ng, 20 ng, 10 ng, 5 ng and 2.5 ng). After blocking with 3% Milk (NON-FAT DRY MILK, Chem-Cruz, Cod SC-2324) in TBS-Tween 0.05% for 30 min, the incubation with the primary antibody anti-Cas9 (MS mAb to CRISPR-Cas9 [7A9-3A3], AbCam) 1μg/ml in blocking solution was performed O/N at 4°C. The membrane was washed 3 times in TBS-Tween 0.05%, and incubated with the secondary antibody Anti-Mouse IgG-Peroxidase (SIGMA-life science, Cod A-9044-2ml, 10-20 mg/mL) in dilution 1:2000 in blocking solution for 30 min at room temperature. After 3 washes in TBS-Tween 0.05% of 5 minutes and one in 1x TBS (20 mM Tris-HCl, 15 mM NaCl, pH 7,5) of 15 minutes, the membrane was developed with ECL chemiluminescent signal (Clarity max™Western ECL Substrate, BioRad, Cod 1705062). The signal was detected by ChemiDoc™ XRS+ Imaging system (BioRad) and the protein quantification was performed in ImageLab TM software (BioRad) using the analysis tool box.

### DNA template synthesis and purification

The selected linear DNA fragment for in vitro experiments contained the first exon of the zebrafish tyr gene (NCBI Reference Sequence: NM_131013.3) and was synthesized by Integrated DNA Technologies (IDT) in a cloning vector (pUCIDT, Amp). We followed a standard protocol for bacterial transformation. Briefly, a 100 μl aliquot of chemically competent bacterial cells (E. Coli, DH5-alpha) was thawed on ice and transferred together with 5 ng of DNA to a prechilled tube. After gentle mixing, the sample was incubated 30 min on ice, followed by heat-shock at 42 °C for 45 sec. The sample was transferred on ice for 2 min, 850 μl of pre-warmed LB Medium was added (SIGMA, Cod. L3522) and incubated at 37 °C on a turning wheel for 45- 60 min. Next, the tube was centrifuged at 12000 x g for 1 min and the ¾ of the supernatant was removed. The pellet was resuspended and spread on a pre-warmed selection plate (Amp, 10 μg/mL, Sigma) and incubated O/N at 37 °C. Obtained single colonies were then incubated O/N at 37°C in 200 mL LB (Amp, 100 μg/mL) broth for plasmid expression. Consequently, plasmid DNA was extracted using the NucleoBond XTRA Midi/ Maxi Kit following the manufacturer’s instructions (Macherey-Nagel, Cod. 7400410.50). Next, standard restriction enzyme digestion by EcoRI (Promega, Cod. 6011) was performed to excise the fragment of interest. The digestion was run on agarose gel as described below (section Agarose Gel Electrophoresis) and the band corresponding to the fragment was excised under UV light. The fragment was purified with the PCR clean up and gel extraction kit according to the supplier’s specifications (Macherey-Nagel, Cod. FC140609L). Fragment concentration was measured by DeNovix DS 11+ FX spectrophotometer/ fluorometer (DeNovix Inc., Wilmington, DE, USA) and stored at – 20 °C until further processing.

### In-vitro cleavage of target DNA

We used the Alt-R CRISPR RNP system (Integrated DNA Technologies,IDT, Coralville, Iowa, USA) to edit our template DNA. The sequence of the gRNA was selected using the CHOPCHOP software. The custom crRNA sequence identified was GGGCCGCAGTATCCTCACTC (+). The RNP complex was assembled according to the protocol suggested for Alt-R-CRISPR Cas9 system-mediated *in vitro* cleavage of target DNA with RNP complex provided on the IDT website. All the previously described components were synthetized by IDT. We varied the RNP concentrations from 0.5 to 1.5μM while maintaining the 50 nM DNA target for all digestion reactions. The mix after the in vitro cleavage assay was loaded in an agarose gel with a final concentration of 1.5% agar prepared in 1x TBE Buffer (89 mM Tris, 89 mM Boric Acid, 2 mM EDTA, pH 8). Ethidium Bromide solution was added for DNA detection (Sigma, Cod. E1510). A 100 bp DNA ladder (Promega, Cod. G210A) was used.

Immediately after completion of the run, the signal was detected using the ChemiDoc™ XRS+ Imaging system (BioRad). ImageLab TM software (BioRad) was used for analysis. Values were normalized to the control and statistical analysis was done and plotted in Graph Pad Prism.

### Zebrafish care

Animal procedures were performed following protocols approved by Italian Ministry of Public Health and the local Ethical Committee of the University of Pisa (authorization n. 99/2012-A, 19.04.2012), in conformity with the Directive 2010/63/EU. Zebrafish were bred in the animal facility of the University of Pisa (Authorization Number DN-16 /43 on 19/01/2015, renewal Authorization Number 1695 on 12/10/2023N). Embryos were obtained by natural mating and maintained at 28 °C in E3 medium (5 mM NaCl, 0.17 mM KCl, 0.33 mM CaCl2, 0.33 mM MgSO4 10-5% Methylene Blue).

### Zebrafish embryo microinjection

Injection solutions were freshly prepared immediately before zebrafish mating started. We used the Alt-R CRISPR-Cas9 System from IDT and adapted the injection protocol by Essner available on the IDT website (Essner J. (2016) Zebrafish embryo microinjection: Ribonucleoprotein delivery using the Alt-R CRISPR-Cas9 System. Coralville, Integrated DNA Technologies. Accessed 21 December, 2017) to our means. Briefly, for the assembly of the RNP we ensured to combine for all experimental setups equimolar amounts of gRNA and Cas9 in the final microinjection volume (6 μL). Any necessary dilutions were performed using the Cas9 dilution buffer (30 mM HEPES, 150 mM KCl, pH 7.5). For better visualization of the injection mix, 0.5 μL of Fast Green dye were added in the 6 μl microinjection volume. As positive control, the RNP was constituted with Cas9 (Alt-R S.p. HiFi Cas9 Nuclease V3). Each zygote was injected in the zygote with 3 nL of microinjection mix using Pneumatic Picopump PV820 air microinjector (World Precision Instruments).

Embryos were checked for any effect of toxicity starting from 5 hours-post-fertilization (hpf), Representative zebrafish embryos images were acquired at 2 or 3 dpf using Nikon stereomicroscope SMZ1500.

### Cell Culture

Human melanoma A375 cells (Cod: ATCC® CRL-1619™) were grown in DMEM (Dulbecco’s Modified Eagle Medium, Cod: 41965-039, GIBCO), 10% FBS (Fetal Bovine Serum, Cod: 10270-106, GIBCO), with addition of GlutaMAX™ supplement (Cod: 35050- 038, GIBCO, 200 mM), and 1x Penicillin/ Streptomycin (Cod: 15140-122, GIBCO) at 37 °C under 5% CO2.

### MTT ASSAY

A375 cells were plated in a 96-multi well plate in complete medium as described above. After 24 h with approximately 80% confluency, cells were treated with different concentrations of nanoparticles in complete medium for 24 h. MTT (3-(4,5-Dimethylthiazol-2-yl)-2,5- Diphenyltetrazolium Bromide, Cod: 5655, Sigma-Aldrich) in dilution 1:10 in complete medium was added in each well. After 1,5 h the medium from each well was removed and 100 μL of DMSO (dimethyl sulfoxide for cell culture, Cas-N°: 67-68-5, EC-N°: 200-664-3, A3672,0250, AppliChem Panreac) were added. The absorbance was measured with a microplate reader (tunable VERSAmax, Molecular Devices) at 570 nm and 690 nm, and the results were analysed in the GraphPad Prism software.

### Transmission Electron Microscopy (TEM) Cells embedding

A375 cells, both treated and control samples, were treated for EM characterization as described elsewhere [29]. Briefly cells were fixed in Petri dishes with 1.5% glutaraldehyde in Na Cacodilate buffer (0.1 M pH 7.4), then scraped, collected in 1.5 ml tubes, and centrifuged for pellet stabilization. After rinses cells were post-fixed in reduced osmium tetroxide (1% K3Fe(CN)6 plus 1% OsO4 in Na Cacodilate buffer 0.1 M pH 7.4). To increase their contrast cells were stained with our homemade X Solution (Moscardini A. et al., 2020), and subsequently dehydrated in a growing series of ethanol concentration. Finally, cells were embedded in epoxy resin (Epon 812, Electron Microscopy Science, Hatfield, PA, USA) and baked for 48h at 60°C.

### Sectioning

Semithin sections (500 nm) were cut using an ultramicrotome (UC7—Leica Microsystems, Vienna, Austria), were collected on glass coverslips and stained with 0.1% ethylene blue, 0.1% toluidine blue in PB. For the ultrastructural analysis, thin 90-nm thick sections were collected on 300 mesh copper grids (G300Cu Electron Microscopy Science, Hatfield, PA, USA) and analyzed with the TEM.

### Confocal microscopy

To evaluate the internalization of the AuNP-Cas9, 20000 cells (either A375 or SK MEL 28) were seeded on a 24 well plate and treated with the nanoformulation 2 days later. After 2 hours of incubation, the cells were fixed in 4% FA for 10 minutes. Then they were washed in PBS 0.1 % Triton and permeabilized in PBS 0.5 % Triton for 15 minutes. Next, they were blocked in PBS 1% BSA and 0.3% Triton for 1 hour and incubated O/N with the primary antibody anti-Cas9 (MS mAb to CRISPR-Cas9 [7A9-3A3], AbCam) 1: 200 dilution in PBS 1% BSA and 0.2% Triton. Next day, the cells were washed 3 times in PBS, and incubated with the secondary antibody Anti-Mouse IgG-Peroxidase (SIGMA-life science, Cod A-9044- 2ml, 10-20 mg/mL) in dilution 1:500, Hoechst 1:750, Phalloidin 1:250 in PBS 1% BSA and 0.2% Triton for 1 hour at room temperature.

### Editing of human melanoma cells

We mixed AuNP-Cas9 and ASAH1 targeting gRNA in an equimolar ratio and incubated for 5 minutes at RT. Then, we seeded 1.2 x 10^5^ A375 melanoma cells in a 96-well plate and administered AuNP-Cas9 in OptiMEM (Thermo Fisher Scientific,Waltham, Massachusetts, 31985062) 24h after. Then, we incubated the cells at 37°C and 5% CO2 for 48 hours, using cells transfected with RNP by lipofection as a positive editing control. Cell genomes were extracted 48h post administration using QuickExtract DNA Extraction Solution 1.0 (LGC Biosearch Technologies, Hoddesdon, United Kingdom, QE09050) following manufacturer’s instructions.

### Digital PCR

We processed the A375 genomes with HindIII restriction enzyme as suggested by the protocol of the ddPCR™ NHEJ Genome Edit Detection Assay used (Bio-Rad, Hercules, California, United States). Then ddPCR was perfomed using a specific custom assay designed using Bio-Rad’s online tool (https://www.bio-rad.com/digital-assays/assays-create/genome). ddPCR reaction and analyses were performed according to previously published protocols [30, [31] using the QIAcuity Digital PCR System (Qiagen, Hilden, Germany).

## Supporting Information

Supporting Information is available from the author. Data and metadata can be downloaded by ZENODO repository (10.5281/zenodo.12701439).

## Acknowledgements

This research was funded by the European Union’s Horizon 2020 Research and Innovation Programme under grant agreement No 862714. Illustrations and experimental schemes were created with Biorender.com.

## References

1. X. Chen, S. Zhong, Y. Zhan, X. Zhang. Crispr-cas9 applications in t cells and adoptive t cell therapies. Cell Mol Biol Lett. 29(1), 52 (2024). 10.1186/s11658-024-00561-1

2. M. L. Zhang, H. B. Li, Y. Jin. Application and perspective of crispr/cas9 genome editing technology in human diseases modeling and gene therapy. Front Genet. 15(1364742 (2024). 10.3389/fgene.2024.1364742

3. M. Jinek, K. Chylinski, I. Fonfara, M. Hauer, J. A. Doudna, E. Charpentier. A programmable dual-rna-guided dna endonuclease in adaptive bacterial immunity. Science. 337(6096), 816-821 (2012). 10.1126/science.1225829

4. M. Jinek, A. East, A. Cheng, S. Lin, E. Ma, J. Doudna. Rna-programmed genome editing in human cells. Elife. 2(e00471 (2013). 10.7554/eLife.00471

5. C. Liu, L. Zhang, H. Liu, K. Cheng. Delivery strategies of the crispr-cas9 gene-editing system for therapeutic applications. J Control Release. 266(17-26 (2017). 10.1016/j.jconrel.2017.09.012

6. M. Behr, J. Zhou, B. Xu, H. Zhang. Delivery of crispr-cas9 therapeutics: Progress and challenges. Acta Pharm Sin B. 11(8), 2150–2171 (2021). 10.1016/j.apsb.2021.05.020

7. R. Luo, H. Le, Q. Wu, C. Gong. Nanoplatform-based in vivo gene delivery systems for cancer therapy. Small. e2312153 (2024). 10.1002/smll.202312153

8. C. Bai, C. Wang, Y. Lu. Novel vectors and administrations for mrna delivery. Small. 19(46), e2303713 (2023). 10.1002/smll.202303713

9. V. Pattanayak, S. Lin, J. P. Guilinger, E. Ma, J. A. Doudna, D. R. Liu. High-throughput profiling of off-target dna cleavage reveals rna-programmed cas9 nuclease specificity. Nat Biotechnol. 31(9), 839–843 (2013). 10.1038/nbt.2673

10. J. Padayachee, M. Singh. Therapeutic applications of crispr/cas9 in breast cancer and delivery potential of gold nanomaterials. Nanobiomedicine (Rij). 7(1849543520983196 (2020). 10.1177/1849543520983196

11. R. Mout, M. Ray, G. Yesilbag Tonga, Y. W. Lee, T. Tay, K. Sasaki, V. M. Rotello. Direct cytosolic delivery of crispr/cas9-ribonucleoprotein for efficient gene editing. ACS Nano. 11(3), 2452–2458 (2017). 10.1021/acsnano.6b07600

12. S. Kim, D. Kim, S. W. Cho, J. Kim, J. S. Kim. Highly efficient rna-guided genome editing in human cells via delivery of purified cas9 ribonucleoproteins. Genome Res. 24(6), 1012–1019 (2014). 10.1101/gr.171322.113

13. T. J. N. Schmidt, B. Berarducci, S. Konstantinidou, V. Raffa. Crispr/cas9 in the era of nanomedicine and synthetic biology. Drug Discov Today. 28(1), 103375 (2022). 10.1016/j.drudis.2022.103375

14. D. Kim, Q. V. Le, Y. Wu, J. Park, Y. K. Oh. Nanovesicle-mediated delivery systems for crispr/cas genome editing. Pharmaceutics. 12(12), (2020). 10.3390/pharmaceutics12121233

15. S. Zhen, Y. Liu, J. Lu, X. Tuo, X. Yang, H. Chen, W. Chen, X. Li. Human papillomavirus oncogene manipulation using clustered regularly interspersed short palindromic repeats/cas9 delivered by ph-sensitive cationic liposomes. Hum Gene Ther. 31(5-6), 309–324 (2020). 10.1089/hum.2019.312

16. Z. Zhang, T. Wan, Y. Chen, H. Sun, T. Cao, Z. Songyang, G. Tang, C. Wu, Y. Ping, F. J. Xu, J. Huang. Cationic polymer-mediated crispr/cas9 plasmid delivery for genome editing. Macromol Rapid Commun. 40(5), e1800068 (2019). 10.1002/marc.201800068

17. B.-C. Zhang, P.-Y. Wu, J.-J. Zou, J.-L. Jiang, R.-R. Zhao, B.-Y. Luo, Y.-Q. Liao, J.-W. Shao. Efficient crispr/cas9 gene-chemo synergistic cancer therapy via a stimuli-responsive chitosan-based nanocomplex elicits anti-tumorigenic pathway effect. Chemical Engineering Journal. 393(124688 (2020). 10.1016/j.cej.2020.124688

18. L. Duan, K. Ouyang, X. Xu, L. Xu, C. Wen, X. Zhou, Z. Qin, Z. Xu, W. Sun, Y. Liang. Nanoparticle delivery of crispr/cas9 for genome editing. Front Genet. 12(673286 (2021). 10.3389/fgene.2021.673286

19. P. Yang, A. Y.-T. Lee, J. Xue, S.-J. Chou, C. Lee, P. Tseng, T. X. Zhang, Y. Zhu, J. Lee, S.-H. Chiou, H.-R. Tseng. Nano-vectors for crispr/cas9-mediated genome editing. Nano Today. 44(101482 (2022). 10.1016/j.nantod.2022.101482

20. G. A. Kanu, J. B. M. Parambath, R. O. Abu Odeh, A. A. Mohamed. Gold nanoparticle-mediated gene therapy. Cancers (Basel). 14(21), (2022). 10.3390/cancers14215366

21. R. Mout, V. M. Rotello. Cytosolic and nuclear delivery of crispr/cas9-ribonucleoprotein for gene editing using arginine functionalized gold nanoparticles. Bio Protoc. 7(20), (2017). 10.21769/BioProtoc.2586

22. K. Lee, M. Conboy, H. M. Park, F. Jiang, H. J. Kim, M. A. Dewitt, V. A. Mackley, K. Chang, A. Rao, C. Skinner, T. Shobha, M. Mehdipour, H. Liu, W. C. Huang, F. Lan, N. L. Bray, S. Li, J. E. Corn, K. Kataoka, J. A. Doudna, I. Conboy, N. Murthy. Nanoparticle delivery of cas9 ribonucleoprotein and donor dna. Nat Biomed Eng. 1(889-901 (2017). 10.1038/s41551-017-0137-2

23. T. Fang, X. Cao, M. Ibnat, G. Chen. Stimuli-responsive nanoformulations for crispr-cas9 genome editing. J Nanobiotechnology. 20(1), 354 (2022). 10.1186/s12951-022-01570-y

24. X. Li, Y. Pan, C. Chen, Y. Gao, X. Liu, K. Yang, X. Luan, D. Zhou, F. Zeng, X. Han, Y. Song. Hypoxia-responsive gene editing to reduce tumor thermal tolerance for mild- photothermal therapy. Angew Chem Int Ed Engl. 60(39), 21200–21204 (2021). 10.1002/anie.202107036

25. E. Ju, T. Li, S. Ramos da Silva, S. J. Gao. Gold nanocluster-mediated efficient delivery of cas9 protein through ph-induced assembly-disassembly for inactivation of virus oncogenes. ACS Appl Mater Interfaces. 11(38), 34717–34724 (2019). 10.1021/acsami.9b12335

26. L. E. Jao, S. R. Wente, W. Chen. Efficient multiplex biallelic zebrafish genome editing using a crispr nuclease system. Proc Natl Acad Sci U S A. 110(34), 13904–13909 (2013). 10.1073/pnas.1308335110

27. Y. C. Wu, I. J. Wang. Heat-shock-induced tyrosinase gene ablation with crispr in zebrafish. Mol Genet Genomics. 295(4), 911–922 (2020). 10.1007/s00438-020-01681-x

28. J. Turkevich, P. C. Stevenson, J. Hillier. A study of the nucleation and growth processes in the synthesis of colloidal gold. Discussions of the Faraday Society. 11(0), 55–75 (1951). 10.1039/DF9511100055

29. A. K. Mapanao, M. Santi, P. Faraci, V. Cappello, D. Cassano, V. Voliani. Endogenously triggerable ultrasmall-in-nano architectures: Targeting assessment on 3d pancreatic carcinoma spheroids. ACS Omega. 3(9), 11796–11801 (2018). 10.1021/acsomega.8b01719

30. S. Crucitta, M. Ruglioni, C. Novi, M. Manganiello, R. Arici, I. Petrini, E. Pardini, F. Cucchiara, F. Marmorino, C. Cremolini, S. Fogli, R. Danesi, M. Del Re. Comparison of digital pcr systems for the analysis of liquid biopsy samples of patients affected by lung and colorectal cancer. Clinica Chimica Acta. 541(117239 (2023). 10.1016/j.cca.2023.117239

31. M. Lai, E. Maori, P. Quaranta, G. Matteoli, F. Maggi, M. Sgarbanti, S. Crucitta, S. Pacini, O. Turriziani, G. Antonelli, L. Heeney Jonathan, G. Freer, M. Pistello. Crispr/cas9 ablation of integrated hiv-1 accumulates proviral dna circles with reformed long terminal repeats. Journal of Virology. 95(23), 10.1128/jvi.01358-01321 (2021).

