## Supplementary data for "A Transfection-Free Approach of Gene Editing via a gold-based nanoformulation of the Cas9 protein"

Supplementary Figures

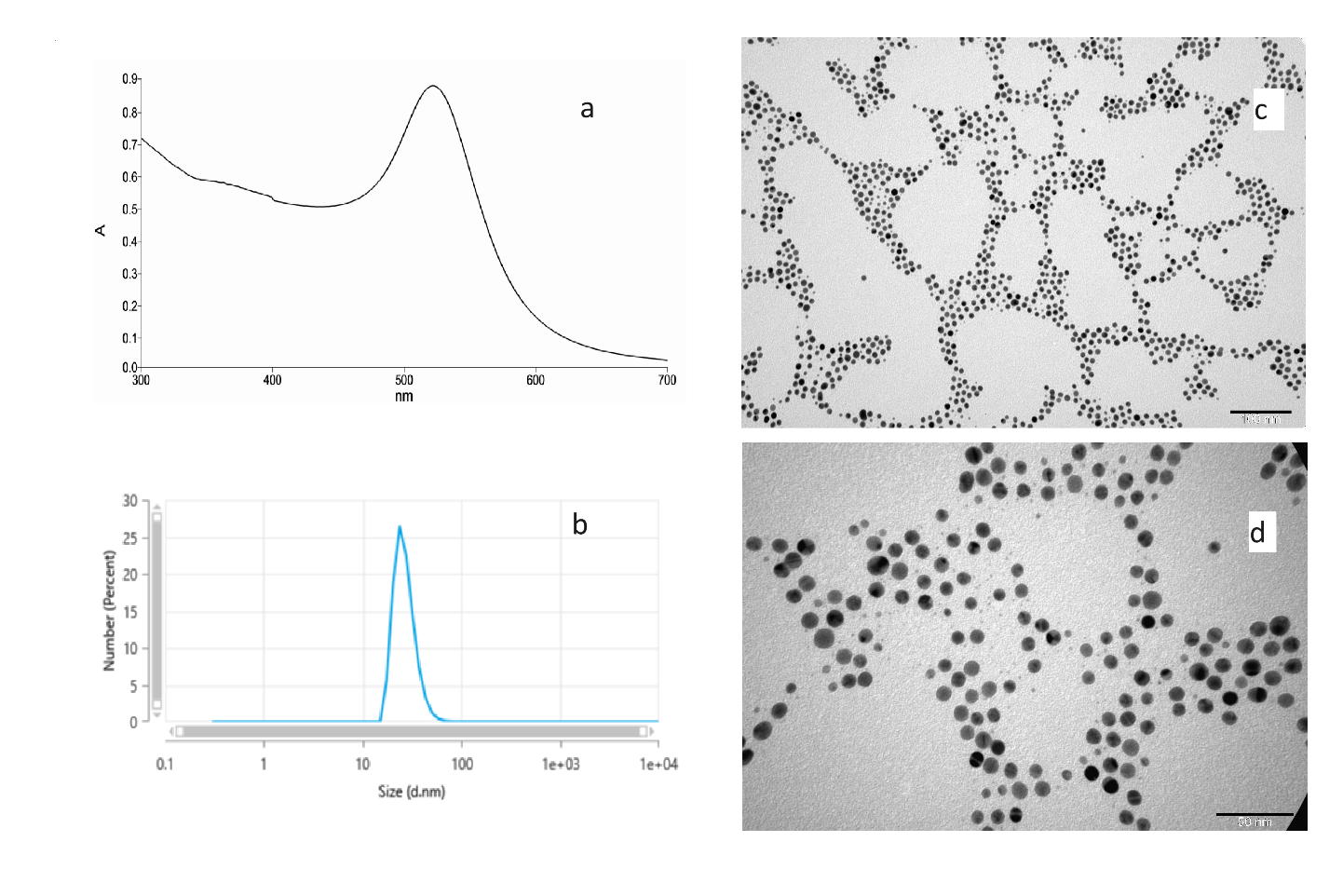

**Figure S1.** Characterization of AuNP with carboxylic background and NTA on the surface. (A) UV-VIS spectra with SPR max 522 ± 3 nm, (B) size distribution (size by DLS 26 ± 6 nm); (C,D) TEM images of metal core 11 ± 3 nm. Zeta potential -35 ± 5mV.

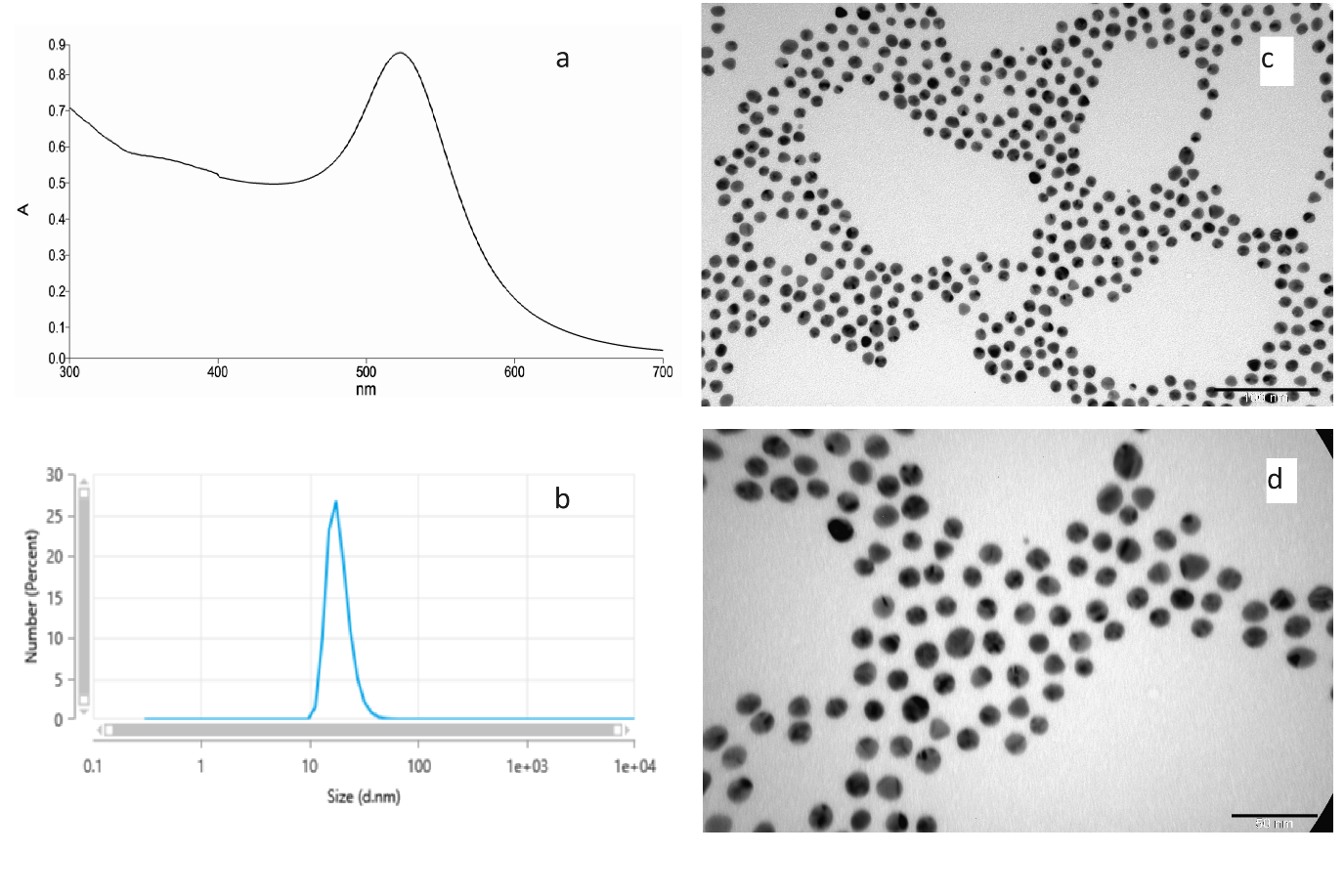

**Figure S2**. Characterization of AuNP with amino background and NTA on the surface. (A) UV-VIS spectra with SPR max 522 ± 3 nm, (B) size distribution (size by DLS 23 ± 5 nm); (C,D) TEM images of metal core 11 ± 3 nm. Zeta potential -2 ± 20mV.

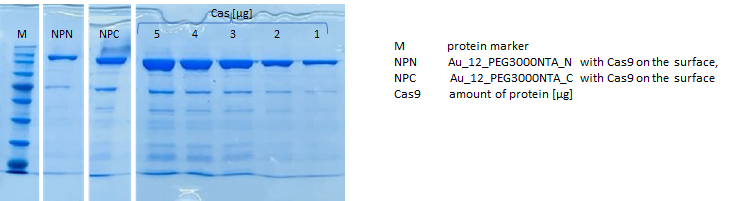

**Figure S3**. Cas9 presence confirmation on the surface of both types of spherical nanoparticles (NPN - AuNP with amino background, NPC - AuNP with carboxylic background).

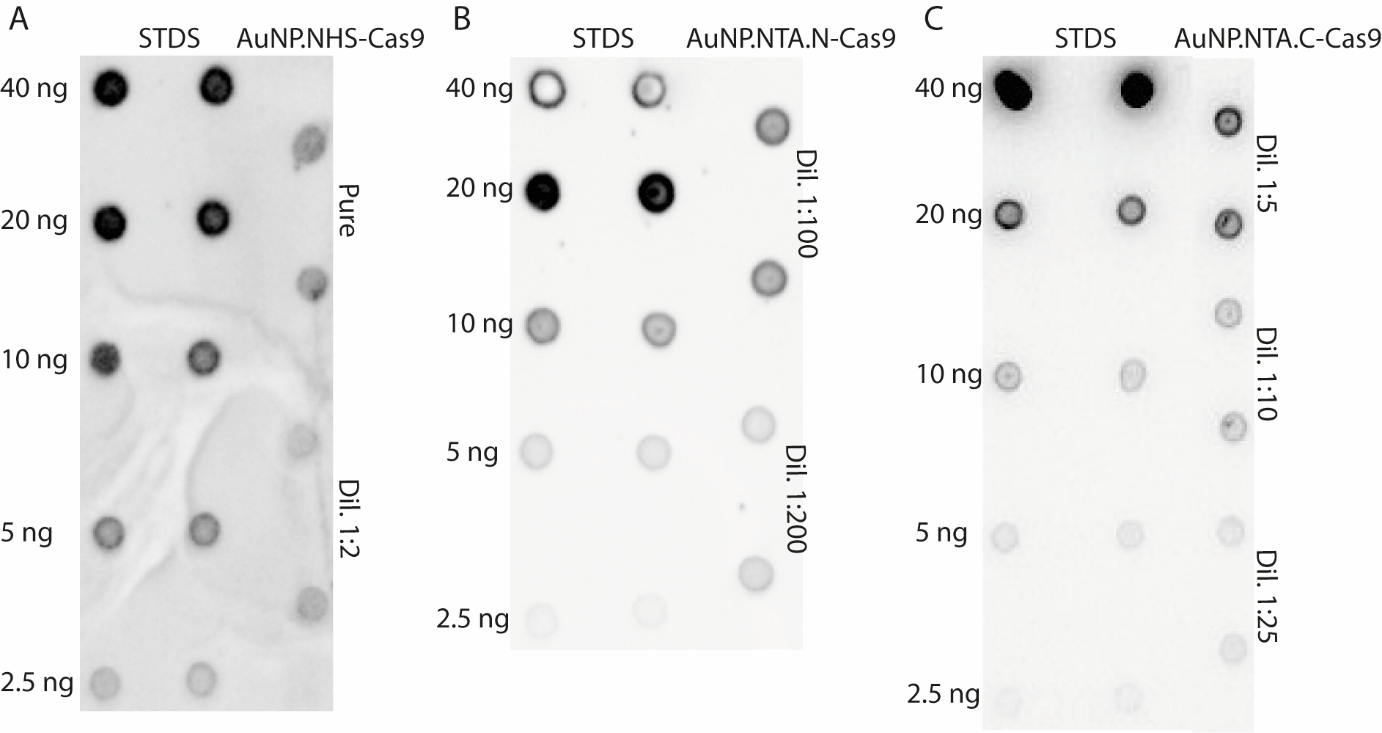

**Figure S4**. Representative Dot blots of AuNP-Cas9 exploiting different spherical nanoparticles; (A) Au_12_NHS, (B) Au_12_NTA.N and (C) Au_12_NTA.C.

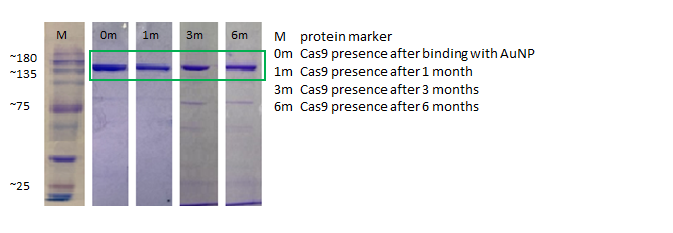
 **Figure S5.** Timeline of Cas9 binding to the surface of AuNP.NTA.C confirmed by SDS-PAGE*.*

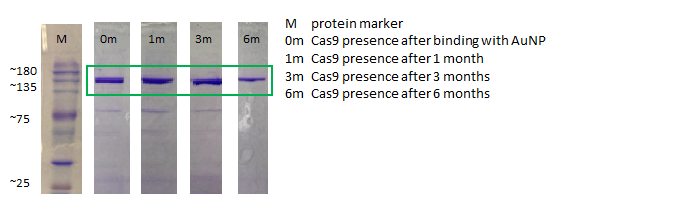

**Figure S6.** Timeline of Cas9 binding to the surface of AuNP.NTA.N confirmed by SDS-PAGE.

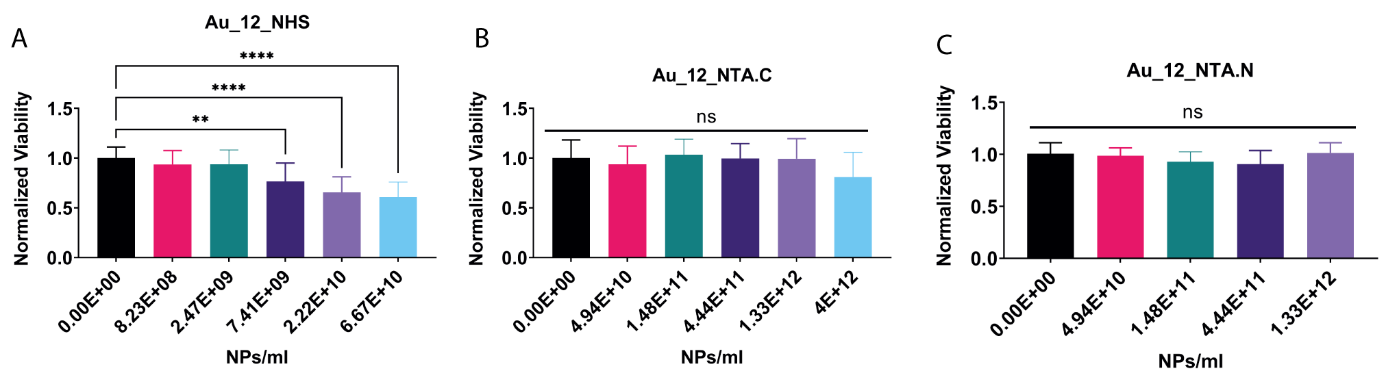

**Figure S7**. Cytotoxicity of AuNPs. Statistical analysis was performed by Kruskal-Wallis test; (A) Au_12_NHS p<0.0001, N=16 per concentration; comparison between 0 and 8.23E+08 and 2.47E+09 p>0.9999, between 0 and 0.7.41E+09 p=0.0048, between 0 and 2.22E+10 and 6.67E+10 p<0.0001, (B) Au_12_NTA.C p=0.4361 N=7 per concentration, (C) Au_12_NTA.N p= 0.1229 N>11 per concentration.

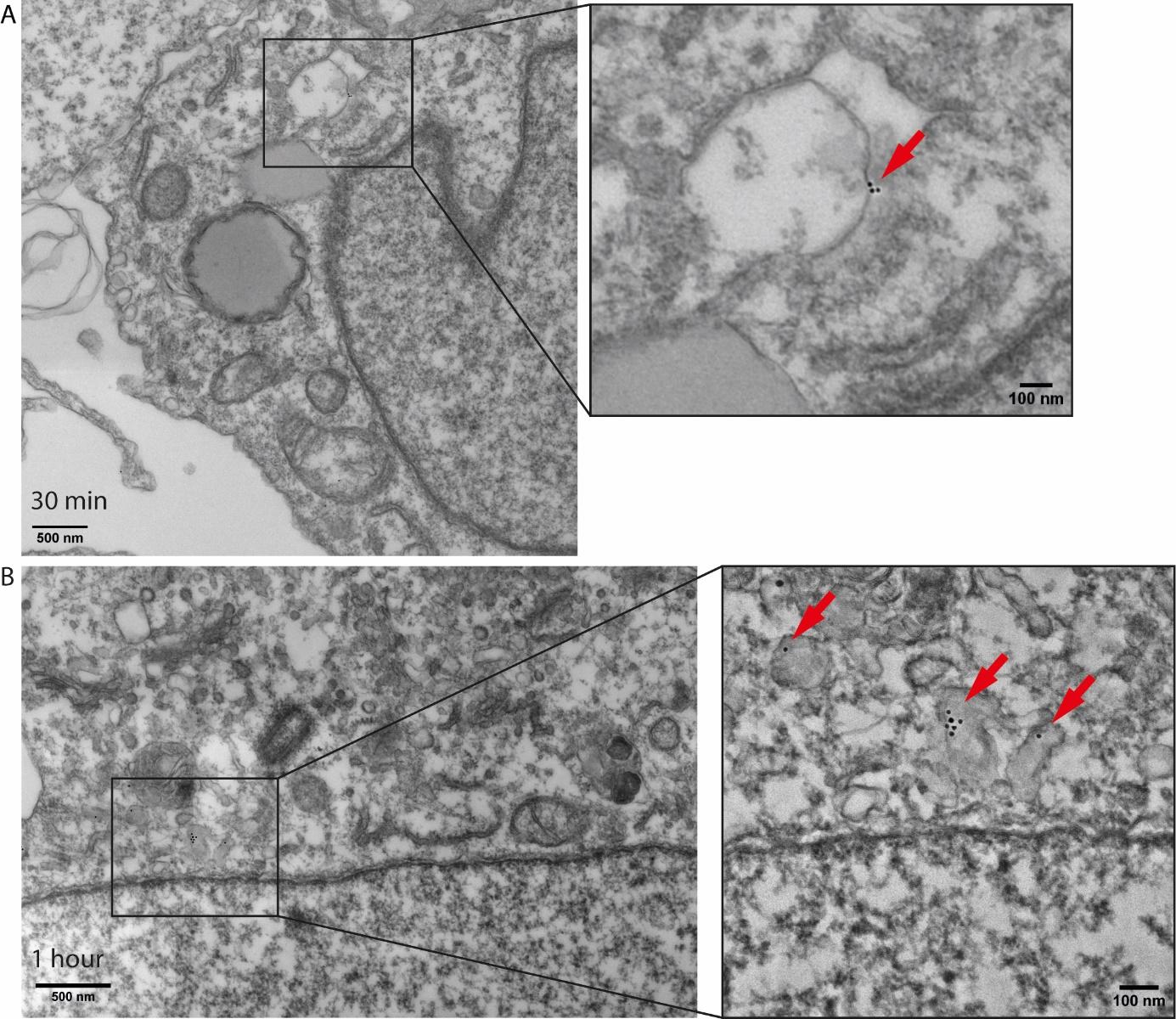

**Figure S8**. TEM images of A375 human melanoma cell line treated with the AuNP.NTA.N-Cas9: gRNA for (A) 30 minutes and (B) 1 hour. The red arrows indicate the AuNP.NTA.N-Cas9: gRNA. The scale bar for the large images is 500 nm and for the magnification 100 nm.

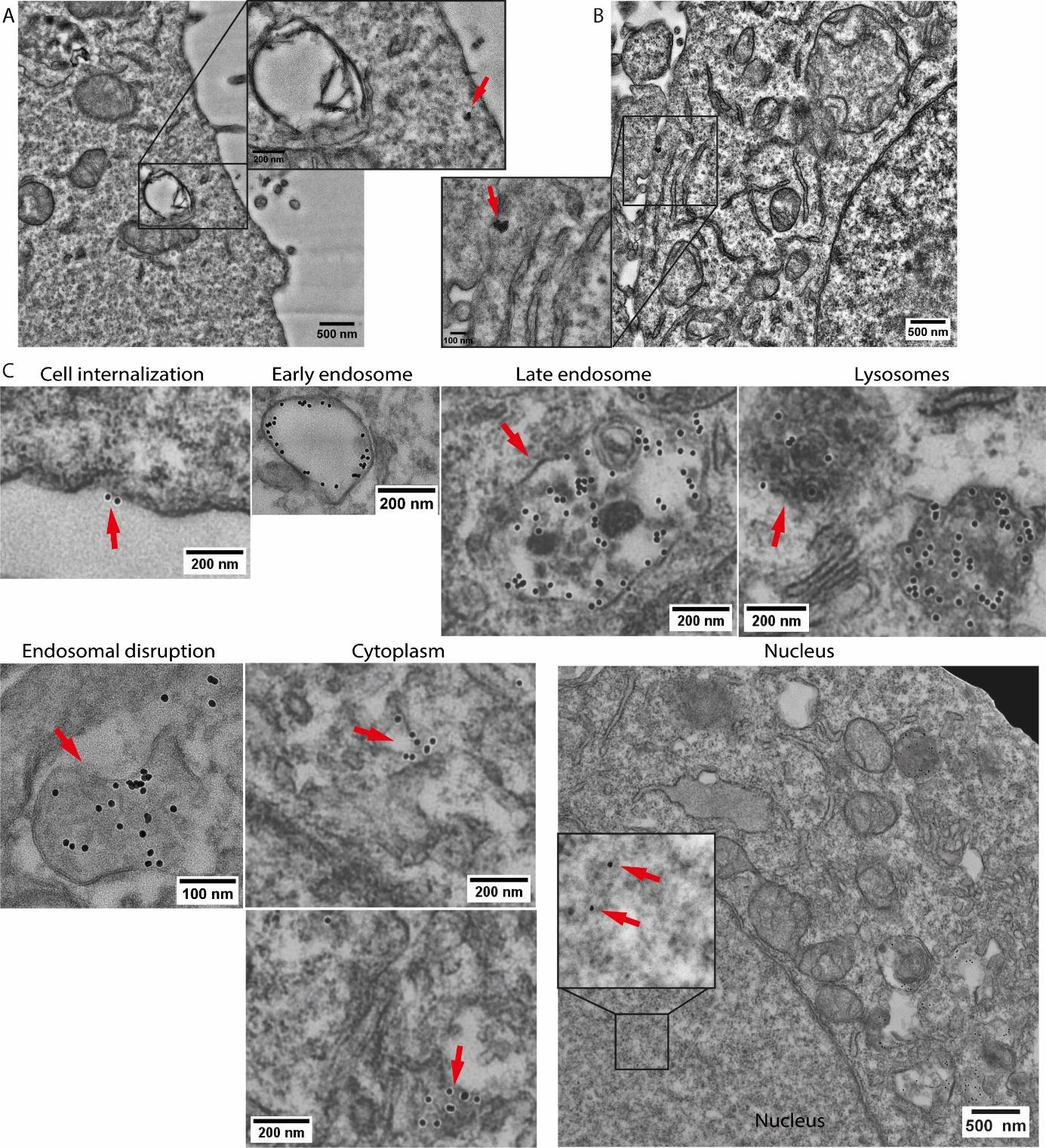

**Figure S9**. TEM images of A375 human melanoma cell line treated with non-functionalized Au_12_NTA.N for 5 hours. The red arrows indicate the naked nanoparticles in agglomerated forms. The scale bar for the large images is 500 nm and for the magnification 200 nm for A, and 100 nm for B.

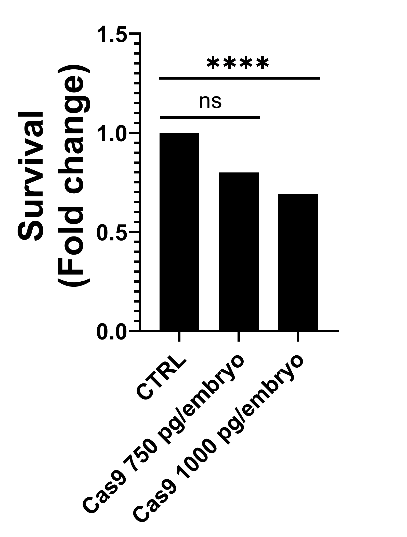

**Figure S10.** Survival index of zebrafish larvae injected with the unconjugated Cas9 in increasing concentrations; 750 pg/embryo and 1000 pg/embryo. Statistical analysis was performed by Chi-square test with p=0.2208, and p<0.0001, respectively.

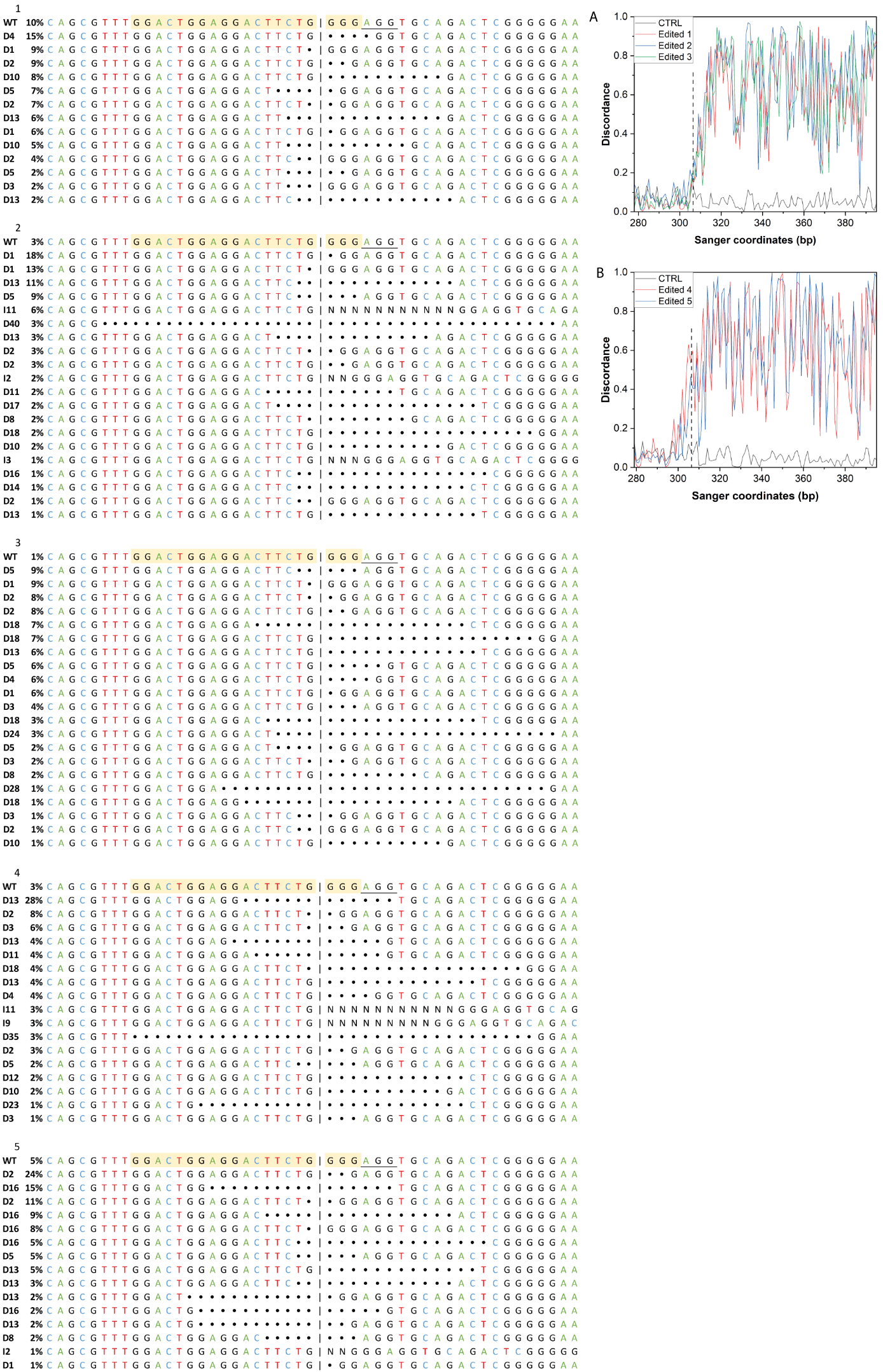

**Figure S11.** Sanger sequencing analysis by ICE software of zebrafish injected with the AuNP-Cas9:gRNA tyr1

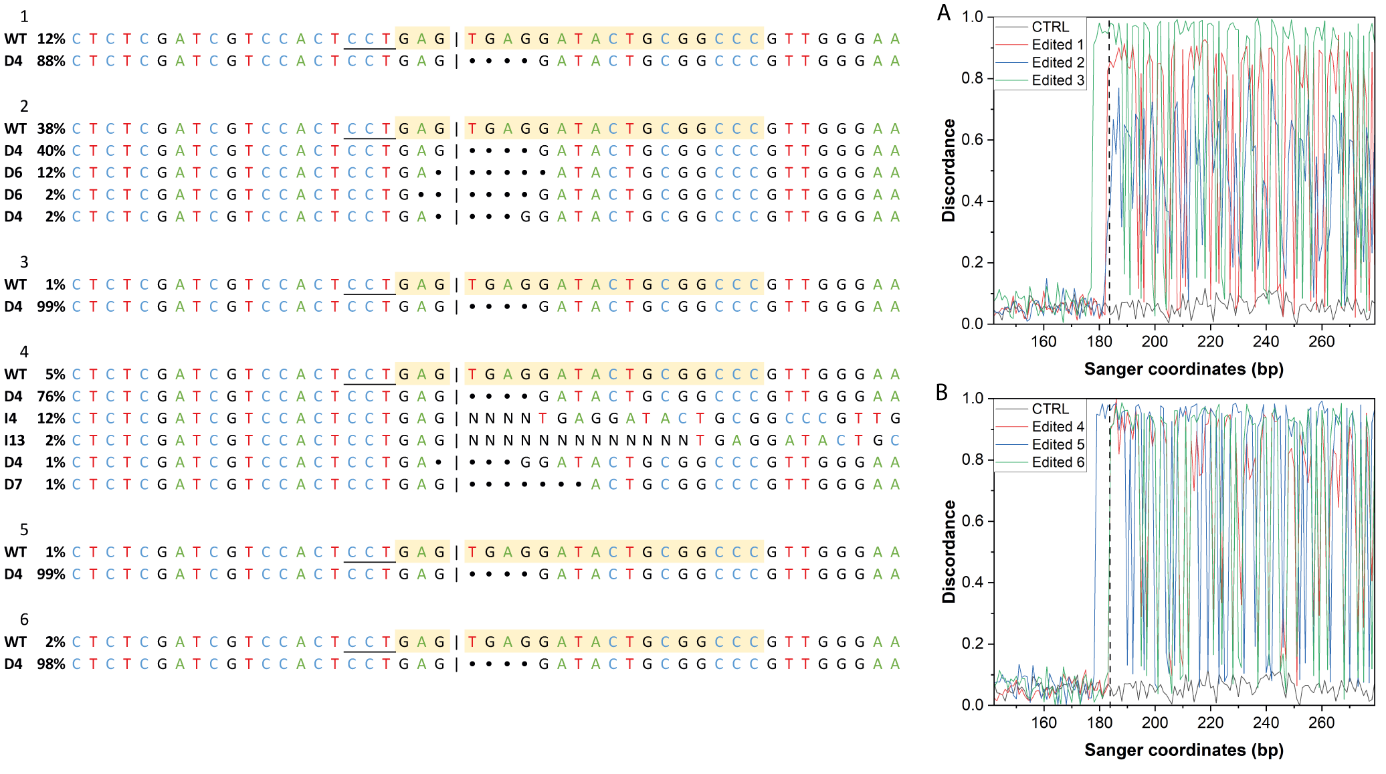

**Figure S12.** Sanger sequencing analysis by ICE software of zebrafish injected with the AuNP-Cas9:gRNA tyr2

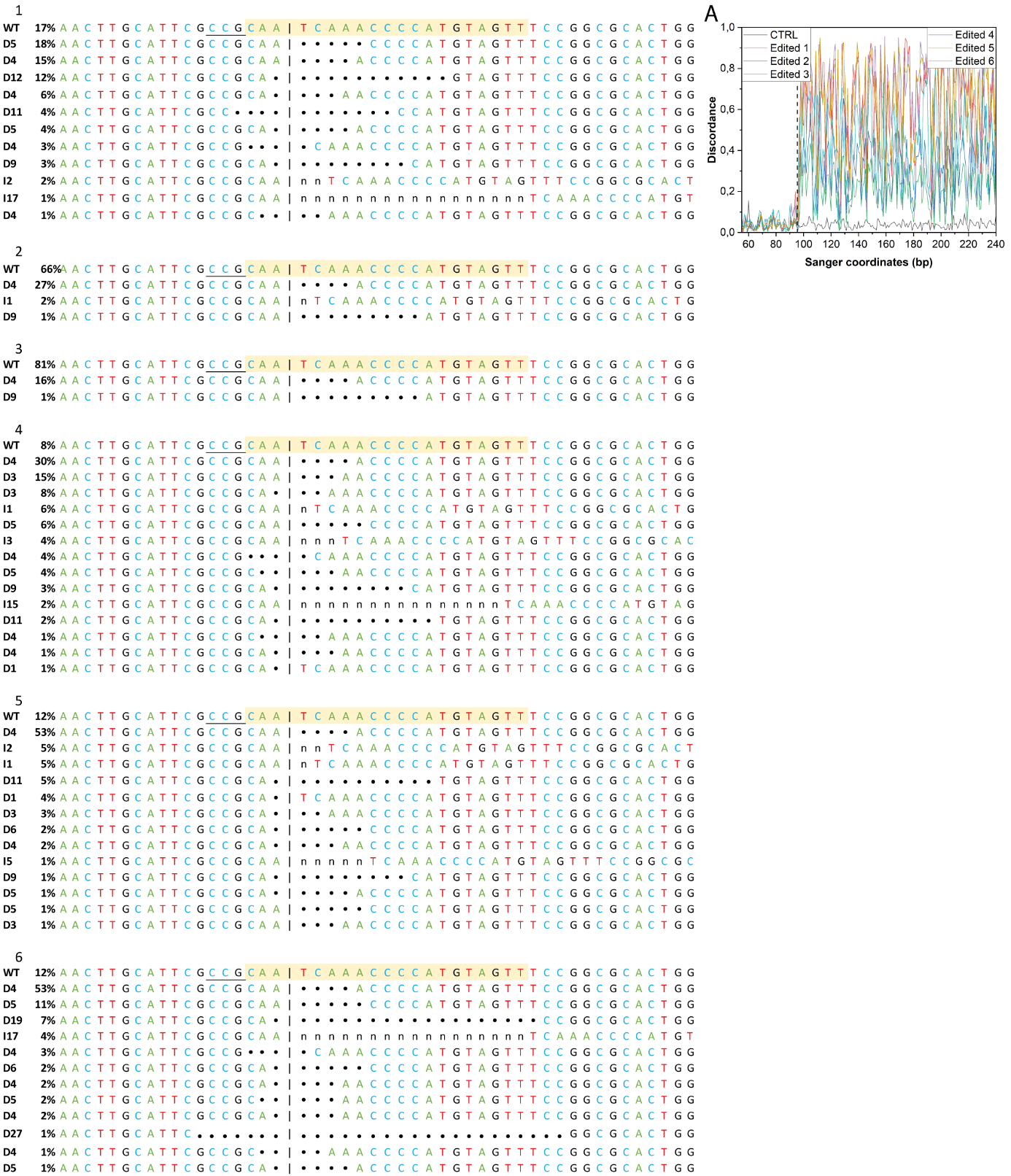

**Figure S13.** Sanger sequencing analysis by ICE software of zebrafish injected with the AuNP-Cas9:gRNA tyr3

Table S1. All functionalization batches tested

|  | **LOT** | **CODE** | **DATE** | **ng Cas9/NP** | **Cas9 molecules/ NP** |
| --- | --- | --- | --- | --- | --- |
| **AuNP.NTA.N-Cas9** | 260122 | F6.NTA.N.Cas9 | 22.03.2022 | 1.59E-08 | 59.1 |
| **AuNP.NTA.N-Cas9** | 260122 | F7.NTA.N.Cas9 | 3.05.2022 | 3.00E-09 | 11.2 |
| **AuNP.NTA.N-Cas9** | 100522 | F8.NTA.N.Cas9 | 14.06.2022 | 9.97E-09 | 37.1 |
| **AuNP.NTA.N-Cas9** | 100522 | F9.NTA.N.Cas9 | 10.08.2022 | 8.57E-09 | 31.9 |
| **AuNP.NTA.N-Cas9** | 100223 | F10.NTA.N.Cas9 | 17.04.2023 | 8.60E-09 | 32.0 |
| **AuNR.NTA.N-Cas9** | 100223 | F11.NTA.N.Cas9 | 17.04.2023 | 5.70E-09 | 21.2 |
| **AuNP.NTA.N-Cas9** | 100223 | F12.NTA.N.Cas9 | 16.05.2023 | 3.17E-08 | 117.8 |
| **AuNP.NTA.N-Cas9** | 150523 | F13.NTA.N.Cas9 | 9.06.2023 | 2.13E-08 | 79.2 |
| **AuNP.NTA.N-Cas9** | 150523 | F14.NTA.N.Cas9 | 9.06.2023 | 9.33E-09 | 34.7 |
| **AuNP.NTA.N-Cas9** | 260923 | F15.NTA.N.Cas9 | 4.11.2023 | 1.60E-08 | 59.5 |
| **AuNP.NTA.N-Cas9** | 112023 | F16.NTA.N.Cas9 | 5.12.2023 | 3.50E-09 | 13.0 |
| **AuNP.NTA.C-Cas9** | 241120 | F1.NTA.C.Cas9 | 9.02.21 | 4.72E-09 | 17.8 |
| **AuNP.NTA.C-Cas9** | 241120 | F2.NTA.C.Cas9 | 17.02.21 | 4.14E-09 | 15.6 |
| **AuNP.NTA.C-Cas9** | 241120 | F3.NTA.C.Cas9 | 22.03.21 | 4.27E-09 | 16.1 |
| **AuNP.NTA.C-Cas9** | 160321 | F4.NTA.C.Cas9 | 28.04.21 | 6.37E-09 | 24.0 |
| **AuNP.NTA.C-Cas9** | 160321 | F5.NTA.C.Cas9 | 31.05.21 | 1.44E-09 | 5.4 |
| **AuNP.NTA.C-Cas9** | 160321 | F6.NTA.C.Cas9 | 04.06.21 | 4.93E-09 | 18.6 |
| **AuNP.NTA.C-Cas9** | 160321 | F7.NTA.C.Cas9 | 04.06.21 | 1.32E-09 | 5.0 |
| **AuNP.NTA.C-Cas9** | 160321 | F9.NTA.C.Cas9 | 18.06.21 | 5.28E-09 | 19.9 |
| **AuNP.NTA.C-Cas9** | 160321 | F10.NTA.C.Cas9 | 18.06.21 | 3.88E-09 | 14.6 |
| **AuNP.NTA.C-Cas9** | 160321 | F11.NTA.C.Cas9 | 13.07.21 | 7.52E-09 | 28.3 |
| **AuNP.NTA.C-Cas9** | 160321 | F12.NTA.C.Cas9 | 13.07.21 | 5.33E-09 | 20.0 |
| **AuNP.NTA.C-Cas9** | 160321 | F13.NTA.C.Cas9 | 27.07.21 | 1.08E-08 | 40.8 |
| **AuNP.NTA.C-Cas9** | 160321 | F14.NTA.C.Cas9 | 01.09.21 | 1.90E-09 | 7.1 |
